# Chlamydomonas cells transition through distinct Fe nutrition stages within 48 h of transfer to Fe-free medium

**DOI:** 10.1101/2024.02.13.579691

**Authors:** Helen W. Liu, Eugen I. Urzica, Sean D. Gallaher, Crysten E. Blaby-Haas, Masakazu Iwai, Stefan Schmollinger, Sabeeha S. Merchant

**Author notes:** ^b^Competence Network IBD, Hopfenstrasse 60, 24103 Kiel, Germany. ^c^California Institute for Quantitative Biosciences (QB3), University of California, Berkeley, CA 94720, USA. ^d^Molecular Foundry, Lawrence Berkeley National Laboratory, Berkeley, CA 94720, USA. ^e^Plant Research Laboratory, Department of Biochemistry and Molecular Biology, Michigan State University, E. Lansing, MI 48824, USA. Address correspondence to Sabeeha S. Merchant, Address: QB3, Stanley Hall, University of California, Berkeley, CA 9720.

## Abstract

Low iron (Fe) bioavailability can limit the biosynthesis of Fe-containing proteins, which are especially abundant in photosynthetic organisms, thus negatively affecting global primary productivity. Understanding cellular coping mechanisms under Fe limitation is therefore of great interest. We surveyed the temporal responses of Chlamydomonas (*Chlamydomonas reinhardtii*) cells transitioning from an Fe-rich to an Fe-free medium to document their short- and long-term adjustments. While slower growth, chlorosis and lower photosynthetic parameters are evident only after one or more days in Fe-free medium, the abundance of some transcripts, such as those for genes encoding transporters and enzymes involved in Fe assimilation, change within minutes, before changes in intracellular Fe content are noticeable, suggestive of a sensitive mechanism for sensing Fe. Promoter reporter constructs indicate a transcriptional component to this immediate primary response. With acetate provided as a source of reduced carbon, transcripts encoding respiratory components are maintained relative to transcripts encoding components of photosynthesis and tetrapyrrole biosynthesis, indicating metabolic prioritization of respiration over photosynthesis. In contrast to the loss of chlorophyll, carotenoid content is maintained under Fe limitation despite a decrease in the transcripts for carotenoid biosynthesis genes, indicating carotenoid stability. These changes occur more slowly, only after the intracellular Fe quota responds, indicating a phased response in Chlamydomonas, involving both primary and secondary responses during acclimation to poor Fe nutrition.

## Introduction

Iron (Fe), in trace amounts, is an essential element for life, used as a crucial cofactor mediating biological redox reactions and reactions involving oxygen chemistry. Although one of the most abundant elements in the Earth’s crust, Fe has limited bioavailability in aerobic environments in typical biology-compatible pH ranges, because Fe is mostly held as insoluble, stable Fe^3+^-oxides (Guerinot and Yi 1994). Photosynthetic organisms, such as phytoplankton and land plants, are particularly affected by the limited Fe bioavailability because of the Fe required for the functioning of their photosynthetic complexes. Indeed, phytoplankton growth in ∼40% of the world’s oceans is Fe-limited (Martin et al. 1994; Boyd et al. 2000; Moore et al. 2001) as is the growth of land plants on 30% of arable land (Chen and Barak 1982), collectively decreasing global primary productivity, and hence exacerbating the potential for food insecurity in face of a growing population and climate change.

In photosynthetic organisms, approximately 40% of Fe is localized to the thylakoid membrane (Raven 1990; Shikanai et al. 2003). Fe is used as a cofactor in all the major membrane protein complexes in oxygenic photosynthesis, comprising of photosystem II (PSII), photosystem I (PSI), the cytochrome (Cyt) *b*_6_*f*, and the soluble electron carriers ferredoxin and, in copper deficiency, Cyt *c*_6_ (Blaby-Haas and Merchant 2004). In response to changes in Fe availability, the stoichiometries of individual photosynthetic complexes are adjusted to optimize photosynthesis (Sherman and Sherman 1983; Sandström et al. 2002). While the overall abundance of the photosynthetic protein complexes decreases in Fe deficiency, PSI is the prime target for degradation because it has the highest Fe content (12 Fe atoms per PSI). Indeed, the ratio of PSI/PSII changes from 2:1 to 1:1 in a cyanobacterium, *Synechococcus*, under Fe deficiency (Sandmann and Malkin 1983). There are two other well-known adjustments to the photosynthetic apparatus in Fe deficiency. One is the replacement of ferredoxin with flavodoxin in which flavin is the redox cofactor instead of a 2Fe-2S center. The replacement, which was initially discovered in bacteria (Ragsdale and Ljungdahl 1984), is widespread in phototrophs, including cyanobacteria, diatoms and some green algae (Pakrasi et al. 1985; Laudenbach et al. 1988; La Roche et al. 1993, 1995; Davidi et al. 2023; Jeffers et al. 2023), although not in the reference alga, *Chlamydomonas reinhardtii* (herein referred to as Chlamydomonas). The second adjustment is the modification of the PSI-associated antenna proteins, presumably to compensate for the lower PSI abundance. This phenomenon is well studied in cyanobacteria where a different light-harvesting complex is associated with PSI in low Fe conditions (Sherman and Sherman 1983; Pakrasi et al. 1985; Boekema et al. 2001; Bibby et al. 2001; Strzepek and Harrison 2004). The second adjustment mechanism may also occur to some degree in algae, but this is not as well studied (Varsano et al. 2006).

Chlamydomonas, which is in the green lineage, is a reference organism widely used for the study of chloroplast metabolism and photosynthesis (Salomé and Merchant 2019). We have developed this alga as a useful system for investigating trace metal homeostasis (Blaby-Haas & Merchant, 2012, 2023; Glaesener et al., 2013; Merchant et al., 2006). It is easy to manipulate the Fe content within the defined growth medium and homogenously supply Fe to cells without variations in organ, tissue, or cell type (Hui et al. 2023). Another strength of Chlamydomonas for metal homeostasis is that nutritional deficiency is possible, in contrast to other systems that rely on chelator-imposed deficiency. Additionally, because of the position of Chlamydomonas’s in the green lineage, any discoveries are also relevant to land plants.

How Chlamydomonas responds to Fe deficiency varies depending on the trophic status of the cells. The alga can grow phototrophically with light and CO_2_, or heterotrophically on acetate as a reduced carbon source, or mixotrophically, which utilizes both CO_2_ and acetate. As a result, the Fe-demanding photosynthetic apparatus is dispensable when acetate is present under low Fe conditions; by contrast, the photosynthetic apparatus is essential and consequently maintained when CO_2_ is the exclusive carbon source (Naumann et al. 2007; Terauchi et al. 2010; Urzica et al. 2012). The loss of photosynthetic complexes occurs by coordinated degradation of the chlorophyll (Chl)–binding protein complexes, starting with the disconnection of the PSI antenna, the degradation of light harvesting complexes I (LHCIs) and PSI, followed by PSII and the Cyt *b*_6_*f* complexes, with light harvesting complexes II (LHCII) retained, possibly as a Chl reservoir (Moseley et al. 2002; Naumann et al. 2007). The abundance of respiratory complexes, which are Fe-dependent, is minimally affected in acetate-grown cells, suggesting that respiration is the preferred metabolic route for production of reducing equivalents and energy in Fe-poor cells (Naumann et al. 2007; Terauchi et al. 2010). These findings speak to strategic metabolic re-prioritization of Fe utilization during Fe insufficiency in the presence of acetate.

Previous studies of Fe nutrition in mixotrophic Chlamydomonas were conducted in defined medium with three distinct stages of growth with respect to Fe nutrition: i) Fe-replete, with 20 µM Fe, which is the standard Fe concentration in a typical Chlamydomonas growth medium and sufficient for maintaining the Fe quota into stationary phase; ii) Fe-deficient, 1–3 µM Fe, where genes involved in high-affinity Fe uptake such as *FOX1* (encoding a multicopper oxidase)*, FTR1* (encoding an Fe permease)*, FRE1* (encoding a ferrireductase), and *FEA1/2* (encoding an extracellular Fe-binding proteins) are induced during log phase growth, although no effect on growth rate is noted; and iii) Fe-limited, less than 0.5 µM Fe supplied, where growth is inhibited so that the culture reaches stationary phase at lower density and photosynthetic protein complexes are reduced (Allen, et al., 2007; La Fontaine et al., 2002; Moseley et al., 2002). Besides the above-mentioned Fe-assimilation proteins, studies have also revealed a previously unknown plastid-localized MnSOD, whose new synthesis increases superoxide dismutase (SOD) activity under poor Fe nutrition, and a candidate Fe transporter, NRAMP4, for intracellular Fe mobilization (Page et al. 2012; Urzica et al. 2012).

In this work, we report on short-term and long-term changes in the Chlamydomonas transcriptome during a transition from Fe-rich to Fe-free medium. Parallel measurements of Chlamydomonas physiology, specifically growth, pigment contents, elemental profiles, and photosynthetic parameters, allow us to contextualize the transcriptome analysis with physiological acclimation events. We further document, through promoter reporter analysis, that transcription of genes encoding Fe-assimilation components is one key regulatory feature of acclimation to poor Fe nutrition in addition to previously demonstrated mechanisms that rely on induced protein degradation for modification of the photosynthetic apparatus (Moseley et al. 2002; Naumann et al. 2005).

## Materials and methods

### Strains and Culture Conditions

All experiments were performed with *Chlamydomonas reinhardtii* strain CC-4532 (wild type, *mt*^−^) or CC-425 (*mt*^+^) for promoter reporter analysis, which are available from the Chlamydomonas Resource Center. Starter cultures were maintained in Tris-acetate phosphate (TAP) medium with trace elements from Hutner’s trace mix for the samples collected for RNA-seq or a revised trace elements for other phenotyping studies (Hutner et al. 1950; Kropat et al. 2011). Cultures were grown at 24°C and 50–100 µmol photons/m^2^/s and shaken continuously at 140 revolutions per minute (RPM). Fe-replete and Fe-depleted states were achieved by maintaining the cells in standard TAP medium (20 µM Fe-EDTA) and, after washing twice with Fe-free TAP medium (containing all trace elements except Fe-EDTA), transferring them to TAP supplemented with or without Fe-EDTA (20 µM Fe).

### Immunoblot Analysis

20 mL of cultures (0.1-2 x 10^7^ cells/mL) were collected at each time point by centrifugation at 2,260x*g* at 4°C for 3 min. Subsequently, we extracted total cell protein by resuspending cell pellets in 300 µL of 10 mM sodium-phosphate pH 7.0. Cells were broken by two cycles of freeze and thaw, where samples were frozen initially in liquid N_2_, thawed slowly at 4°C, and then re-frozen slowly at −80°C and finally re-thawed at 4°C before determination of protein concentration with a Pierce BCA assay against bovine serum albumin (BSA) as a standard (Thermo Fisher Scientific). Proteins were separated by denaturing SDS-PAGE (10 - 15% (w/v) acrylamide monomer) with 10 µg of protein per lane for 1 h at 160 V (Hoefer – Mighty Small II) and transferred to a 0.1 µm nitrocellulose membrane (Amersham Biosciences Protran) by semi-dry electroblotting for 1 h under constant current (60 mA) (Thermo Fisher Scientific) in filter paper soaked in transfer buffer (assembly order from the cathode to the anode: 1) filter paper soaked in T1 buffer: 0.025 M Tris-HCl [pH 10.4] with 0.06 mM ε-aminopropanoic acid, 20% (w/v) isopropanol; 2) gel; 3) nitrocellulose membrane; 4) filter paper soaked in T2 buffer: 0.025 M Tris-HCl [pH=10.4], 20% (w/v) isopropanol; 5) filter paper soaked in T3 buffer: 0.3 M Tris-HCl [pH 10.4], 20% (w/v) isopropanol). After blocking in 3% (w/v) nonfat dried milk in 1x phosphate buffered saline (PBS; 137 mM NaCl, 2.7 mM KCl, 10 mM Na_2_HPO_4_, 1.8 mM KH_2_PO_4_) with 0.1% (w/v) Tween 20 (PBS-T) for 1 h at room temperature, membranes were incubated overnight at 4°C in the following primary antibodies used at the indicated dilutions in the same solution: ferroxidase (FOX1) 1:500 (La Fontaine et al. 2002; Agrisera AB ASO6 120), Cyt *f* 1:1,000 (Xie and Merchant 1996; Agrisera AB AS06 119) and CF_1_ α/β 1:50,000 (Merchant and Selman 1983, Agrisera AB AS03 030). The membranes were subsequently washed once for 15 min, and then washed three additional times at 5 min intervals in PBS-T. Washed membranes were incubated in a 1:6000 dilution of goat anti-rabbit secondary antibody (Southern Biotech) conjugated to alkaline phosphatase in 3% (w/v) nonfat dried milk in PBS-T. The membranes were subsequently washed again for 15 min, and then washed three times at 5 min intervals PBS-T. For visualization of bound antibody, washed membranes were incubated for 0.5–1 min with 10 mL alkaline phosphatase buffer (100 mM Tris-HCl [pH 9.5], 100 mM NaCl, 5 mM MgCl_2_), 0.006% (w/v) nitro blue tetrazolium (NBT), and 0.003% (w/v) of 5-bromo-4-chloro-3-indolyl phosphate *p*-tolidine salt (BCIP).

### Chl Content Determination

Chl was extracted from whole cells in an 80% acetone 20%methanol (v/v) mixture. The samples were centrifugated at 21,130x*g* for 5 min at 25 °C before the absorbance of the supernatant was measured at 647 nm and 664 nm according to (Porra et al. 1989).

### Intracellular Metal Content Determination

Intracellular metal and sulfur (S) contents were determined by ICP-MS/MS as described (Schmollinger et al. 2021) with minor modifications. Briefly, Chlamydomonas cultures at the indicated times throughout the time course were collected by centrifugation at 2,260x*g* for 3 min in a 50 mL Falcon tube at 25°C. The cells were washed twice in 1 mM Na_2_-EDTA at pH 8 to remove cell surface-associated metals. Cells were resuspended in 10 mL Milli-Q H_2_O for a final wash to remove Na_2_-EDTA and collected by centrifugation in a 15 mL Falcon tube. The cell pellet was overlaid with 143 µL of 70% nitric acid (trace metal grade, A467-500, Fisher Scientific) and incubated at 65°C for 16 h before dilution with 9.5 mL Milli-Q H_2_O to a final nitric acid concentration of 2% (v/v) with Milli-Q water. Metal and S contents were determined on an Agilent 8900 Triple Quadrupole ICP-MS/MS instrument, against an environmental calibration standard (Agilent 5183-4688), a S (Inorganic Venture CGS1) and P (Inorganic Ventures CGP1) standard, using ^89^Y as an internal standard (Inorganic Ventures MSY-100PPM). The levels of all analytes were determined in MS/MS mode, where ^56^Fe were directly determined using H_2_ as a cell gas, while ^32^S were determined via mass shift from 32 to 48 utilizing O_2_ in the collision/reaction cell. An average of four to five technical replicate measurements was used for each individual sample. The average variation in between the technical replicate measurements was below 1.8% for all individual experiments and never exceeded 5% for any individual sample.

### Nucleic Acid Analysis

Total RNA was extracted from Chlamydomonas cells as described previously (Quinn and Merchant 1998). RNA quality was assessed on an Agilent 2100 bioanalyzer and by RNA blot hybridization as described previously (Allen, et al., 2007). The probe used for detection, *CBLP* (also reported as *RACK1*), is a 915 bp *EcoRI* fragment from the cDNA cloned in *pcf8-13* (Schloss 1990). For the RNA-seq experiment, duplicate cDNA libraries were prepared from 4 µg of total RNA for each sample in the 0–48 h time course using an Illumina TruSeq RNA Sample Preparation kit version 1. Indexed libraries were pooled and sequenced on an Illumina HiSeq 2000 instrument, with three libraries per lane, as 100 bp single end reads. Raw and processed sequence files are available at the NCBI Gene Expression Omnibus (accession number GSE44611).

Reads were mapped to the Chlamydomonas reference genome (v5 assembly, v5.5 annotation, available from https://Phytozome.jgi.doe.gov) with RNA STAR (Dobin et al. 2013) with --outFilterMismatchNoverLmax 0.04 --alignIntronMax 10000 --outFilterType BySJout -- outSAMstrandField intronMotif. Normalized transcript abundances were calculated in terms of fragments per kb of transcript per million mapped reads (FPKMs) with cuffdiff (v2.0.2) with --multi-read-correct --max-bundle-frags 1000000000. FPKM values were computed from the average expression levels of two independent cultures from genes with expression estimates of at least 1 FPKM at any time point. Differential expression analysis was performed using the DESeq2 package in R (Love et al. 2014). *P*-values obtained from DESeq2 were adjusted for multiple testing using Bejamini-Hochberg correction to control for false discovery rates. Gene ontology (GO) enrichment analysis using the R package topGO 2.40.0 (Alexa et al. 2006) and the GO annotation table from (Lin et al. 2022).

### Promoter Reporter Constructs

Promoter fusion constructs were generated as described in (Blaby and Blaby-Haas 2018). Primers were designed using the Chlamydomonas genome (v4 assembly, available from https://mycocosm.jgi.doe.gov/mycocosm/home) are listed in Table 1. The resulting plasmids were linearized with *BsaI* for *FRE1* or *PsiI* for *FEA2*, *IRT1* and *NRAMP4* and used to transform CC-425 by electroporation together with linearized pARG7.8 as described in (Blaby and Blaby-Haas 2018). Colonies representing Arg prototrophs were grown for 23 days after which each colony was inoculated into a well of a 96-well microplate containing 200 µL TAP per well. After 6 days, each transformant culture was tested by PCR for the presence of the co-transformed reporter construct, and 10 µL was used to inoculate a well in a fresh microplate containing either 200 µL of TAP or TAP minus Fe. The microplate cultures were grown for another 6 days at which point arylsulfatase activity was assayed using α-naphthyl sulfate potassium salt as described in (Blaby and Blaby-Haas 2018) except the absorbance at 665 nm was used instead of 750 nm for normalization.

**Table 1.**
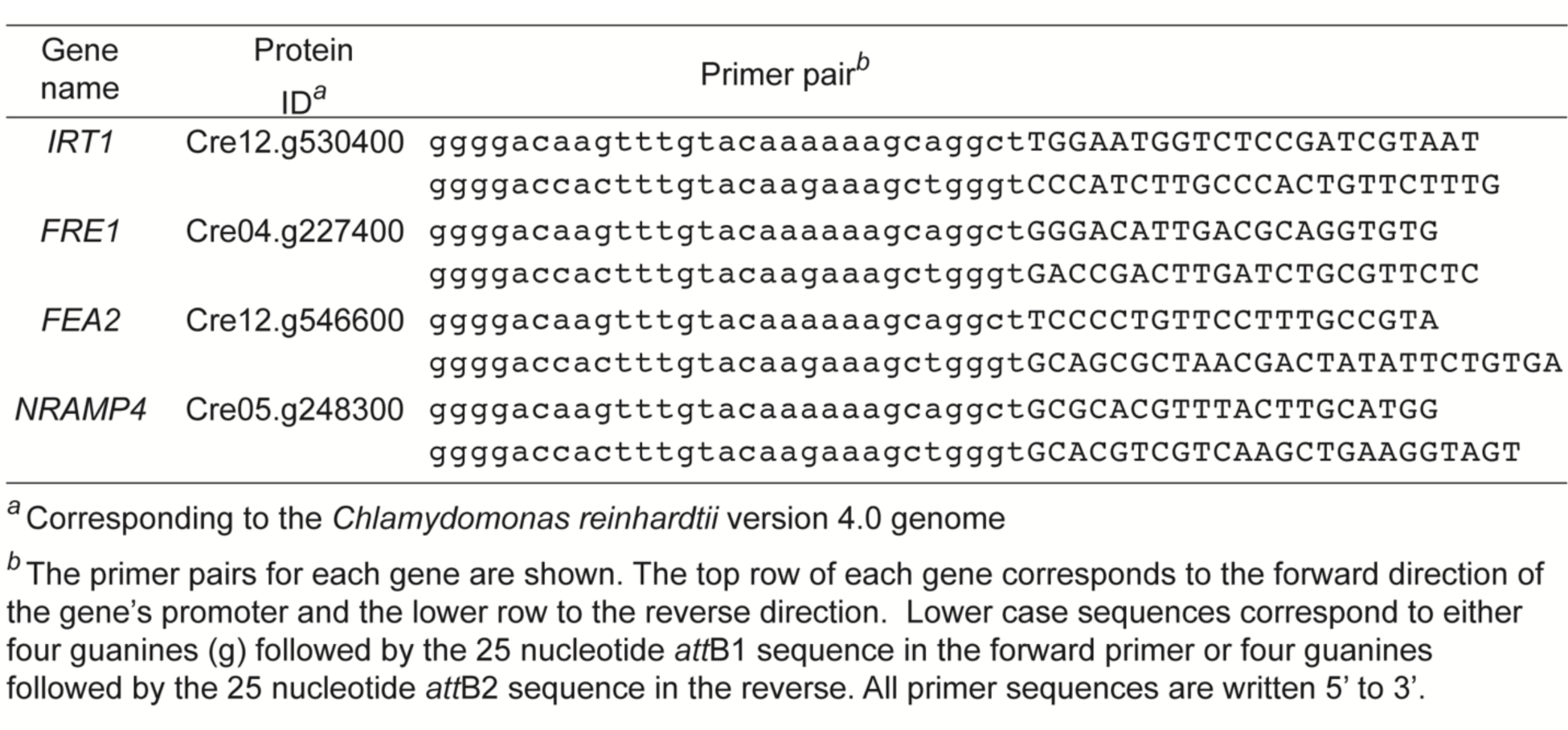
Primers used for reporter constructs.

### High-performance Liquid Chromatography (HPLC) Analysis of Pigments

We collected 10 mL of cultures by centrifugation at 2,260x*g* at 4°C for 3 min either prior to (0’ h) or after transfer of cells to fresh medium (0, 24, 48 h). The pellets were quickly frozen in liquid N_2_. The cells were thawed at room temperature, and the pigments were extracted with 100 µL of 100% (v/v) acetone by vortexing for 10 min. Cell debris was removed by centrifugation at ∼21,000x*g* for 5 min at 4°C, and the supernatant was transferred to a new tube. The remaining pigments in the pellet were extracted a second time with 100–400 µL of 100% (v/v) acetone in the same way described above. The two supernatants were pooled. Extracted pigments were analyzed by using a Spherisorb 5-µm ODS1 column (Waters Corp) according to the method described in (Müller-Moulé et al. 2002).

### Chl Fluorescence

Chl fluorescence (QY_max_) was measured using an AquaPen-C (AP 110-C, Photon Systems Instruments). Cells were diluted to 1 x 10^6^ cells/mL to the final volume of 3 mL and dark-acclimated for 15 min. Cells were exposed to a saturating pulse of ∼400 µmol photons/m^2^/s to probe the photosynthetic parameters of the cells. Chl fluorescence parameters were assayed and calculated according to the definitions of (Baker 2008).

### Accession Numbers

All sequencing data have been deposited at the US National Center for Biotechnology Information Gene Expression Omnibus database under accession number GSE44611.

## Results

### Transfer to Fe-free medium generates Fe limitation within 48 h with a temporal sequence of events

Previously, when we monitored the abundance of various chloroplast-localized Fe-containing proteins after transferring mixotrophic Chlamydomonas cells from Fe-replete to fresh Fe-free TAP medium, we noted dramatic decreases in the abundance of Fe superoxide dismutase (FeSOD), ferredoxin, and Cyt *f* in the first 24 h, but no change in the cellular growth rate despite their lower Fe content (Page et al. 2012). Components of Fe assimilation were already highly induced at that stage, as evidenced by the abundance of the ferroxidase involved in high-affinity Fe uptake and plastid ferritin (J. C. Chen et al., 2008; J. C. Long & Merchant, 2008 and see below). The situation is reminiscent of the Fe-deficient state (Glaesener et al., 2013, and see Introduction). By the second day in Fe-free conditions, ferroxidase accumulation was further increased, and the cells showed clear growth inhibition, reminiscent of the Fe-limited state. This observation suggests that as cells transition from Fe-replete to Fe-poor situations, they experience definable physiological states that we previously designated as Fe deficient and Fe limited (Moseley et al. 2002; Glaesener et al. 2013).

To monitor the physiology of the transition from the Fe-replete to the Fe-deficient to the Fe-limited states and the temporal order of events, we undertook a time course experiment over a 48 h period with dense sampling in the early time points as indicated (Fig.1a). Fe-replete (20 µM) mixotrophic cells at ∼4 x 10^6^ cells/mL (labeled 0’) were collected, washed twice in Fe-free TAP medium and resuspended in fresh medium either supplemented with Fe (labeled 20 µM) or not (labeled 0 µM) to a final density of 2 x 10^6^ cells/mL (time point 0 h). Samples were collected for analysis prior to transfer to new medium (0’ h) or 0 to 48 h after transfer to new medium to assess growth, Fe content and abundance of sentinel proteins for Fe status (Fig.1a).

**Fig.1.**
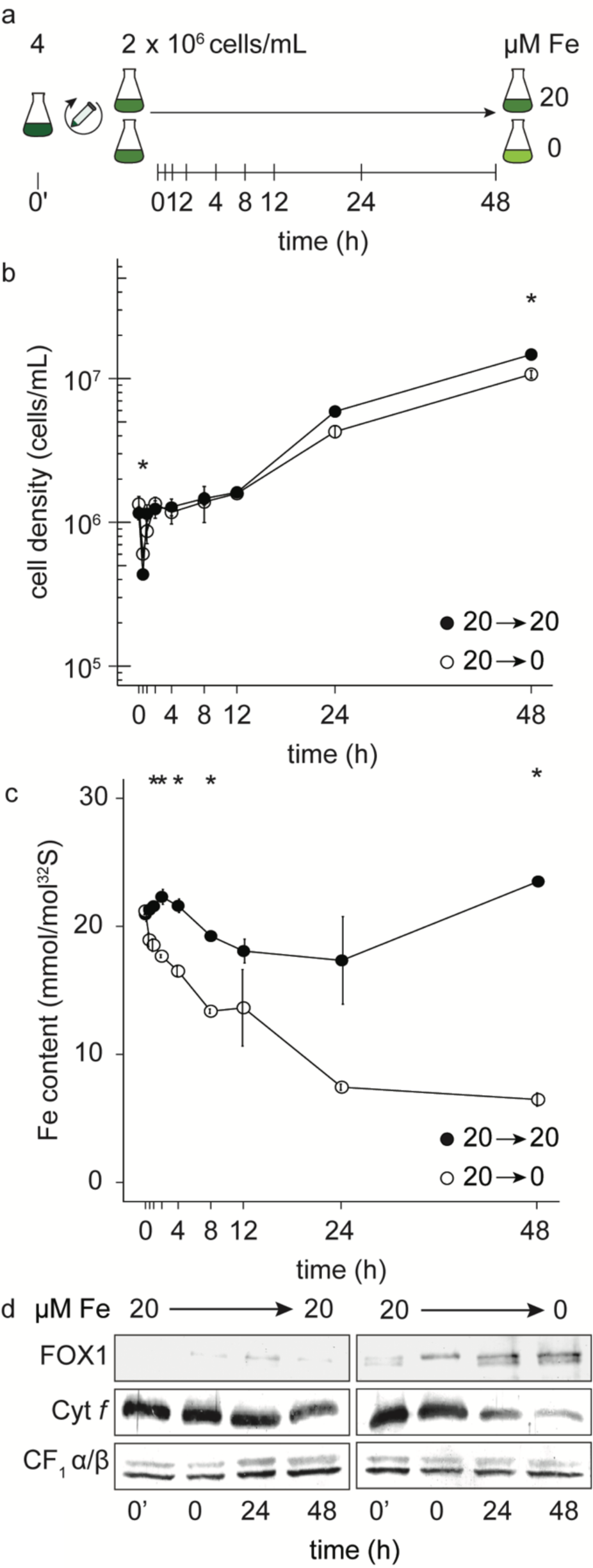
Growth is impaired in Fe-free medium within 48h. (a) Schematic overview of the experiment, (b) cell density and (c) Fe content of Chlamydomonas cells at time point 0’ from Fe-replete medium transitioned into either 20 μM Fe (filled-circle) or 0 μM Fe (open-circle) TAP medium. Vertical lines in (a) indicate sampling times (sampling times under 1 h are not labelled). Sampling procedures are described in the Materials and Methods. Fe content was determined by ICP-MS/MS and normalized to ^32^S content. Error bars indicate the standard error of three independent cultures. Asterisks indicate significant differences between cells in 20 versus 0 μM Fe media at the corresponding sampling times (Student’s *t*-test, *p* ≤ 0.05). (d) Abundance of protein markers for Fe nutrition in Chlamydomonas total cell lysates. 10 µg of protein was separated by denaturing gel electrophoresis and transferred to nitrocellulose for immunoblot analysis. The abundances of ferroxidase (FOX1) and Cyt *f* was monitored using specific antisera, ATP synthase α/β subunits (CF_1_) served as a loading control.

Fe-replete cells that were transferred into fresh Fe-replete medium were able to maintain growth throughout the time course (Fig.1b). Nevertheless, their intracellular Fe content, which initially increased by 8% at 2 h, decreased subsequently at 4 h and remained constant until 24 h despite the high Fe content of the fresh medium (Fig.1c). This observation is consistent with previous work noting this transient decrease in Fe content per cell during rapid exponential growth, because of the inability of Fe assimilation to keep up with biomass accumulation (Page et al. 2012). In contrast, when Fe-replete cells are transferred to Fe-free medium, their Fe content decreased significantly within 30 min, dropping to ∼50% of the Fe content of the Fe-replete control in the first 24 h with minimal effect on growth rate (Fig.1b, c). Although growth rate is not significantly impacted, within the first 24 h, the cell display symptoms of poor Fe nutrition, as evidenced by the decrease in Chl content (Fig.S1) and ferroxidase accumulation (Fig.1d). Within another 24 h, Cyt *f* abundance decreases (Fig.1d) with Chl content remaining at about 50% relative to the Fe-replete cells, consistent with previous results on fully acclimated Fe-limited cultures (Moseley et al. 2002; Terauchi et al. 2010; Devadasu et al. 2016). Fe limitation at 48 h is evident with decreased biomass in the Fe-free culture compared to the Fe-supplemented culture (Fig.1b). The above physiological parameters indicate a continuous temporal progression from Fe-replete through an Fe-deficient to an Fe-limited state within 48 h.

### Long distance view of changes in mRNA abundance during the transition from Fe replete to Fe deplete

RNA was prepared from two separate time course experiments where cells were collected after transfer from Fe-replete to Fe-free medium: a short time course (0 to 4 h) to capture immediate and early responses to the change in Fe status; and a long time course (0 to 48 h) to capture acclimation events and the sustained acclimated state (Fig.2a). Transcript abundances were analyzed by RNA-seq on an Illumina platform (see Methods). RNAs corresponding to 13,028 and 13,770 genes were quantified in the short and long time course experiments, respectively. Five time points, 0, 0.5, 1, 2 and 4 h, were replicated in both experiments to enable comparisons. A principal component analysis (PCA) showed that 64% of the variance in the expression estimates is captured by the first two components, with time in Fe deficiency being the driver as PC1 (Fig.2b). The five time points common to both experiments could be grouped together (compare triangles and circles), indicating consistency between experiments (Fig.2b). Within the PCA, we grouped the samples by amount of time spent in Fe deficiency as follows: 1) the Fe-replete group, comprised the Fe-replete samples and all samples up to the first 15 minutes after transfer to Fe-free medium (0, 5, 10 or 15 min), 2) the early transition group, with samples between 0.5 h and 4 h after transfer in both time courses, and 3) the acclimated state group, corresponding to the Fe-limited state, with all the later time points.

**Fig. 2.**
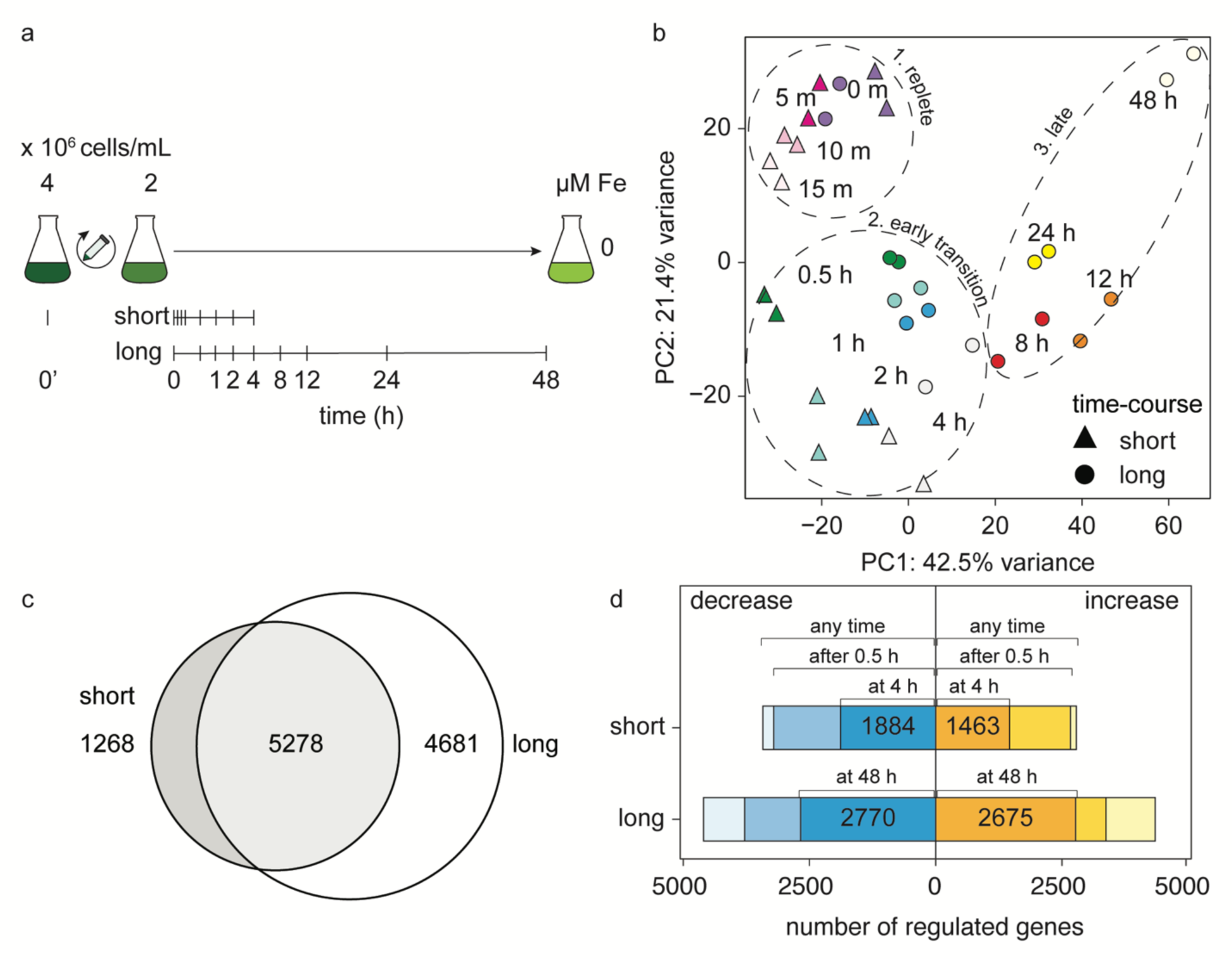
Time without Fe influences mRNA abundances. (a) Schematic overview of the Fe limitation time course. Vertical lines indicate sampling times (in the short and long time courses). The sampling procedures are as described in the Materials and Methods. (b) Principal component analysis of samples collected in the short time course (triangles) and samples collected for the long time course (circles). Each point corresponds to the length of time in Fe-omitted medium. The percentage of the total variance accounted for by the first and second principal components are indicated on the axes. Colors indicate the three phases of transition into Fe-free medium: 1. replete phase, 2. early transition phase, and 3. late phase. The lighter colors represent the later time points within each phase. (c) The intersect of the differentially accumulated transcripts in the short and long time courses (exclusively differentially expressed in short = dark gray, 1,268 genes; shared = light gray, 5,278 genes; and only differentially expressed in long = white, 4,681 genes). (d) Increased (yellow) or decreased (blue) mRNA abundances that were either still changed in the last time point (4 h for the short time course; 48 h in the long time course), changed by 30 min, or changed at any time point. (top) 6,207 DEGs in the short time-course and (bottom) 8,945 DEGs in the long time-course that were exclusively upregulated or downregulated. Genes included in the analysis had a minimal expression of 1 FPKM in at least one time-point in the experiment, experienced a 2-fold change, and a Benjamini-Hochberg-adjusted *p* value ≤ 0.01 for differential expression.

A summary of the differential expression analysis compared to the 0 h time point is shown in Fig. 2c,d (Supplemental Dataset S1). A substantial proportion of the transcriptome, 6,546 genes (∼50% of expressed genes), was differentially expressed in the short time course for at least one time point during the experiment. Of these, mRNA abundances for 2,785 genes increased, while mRNA abundances for 3,422 genes decreased (Fig.2d); a small fraction, 339 genes, showed a pattern of increased mRNA abundances at one stage but decreased abundances at another stage during the time course (Fig.S2, Supplemental Dataset S1). Most (96% up and 94% down) of the changes in the short time course occurred between 30 min and 240 min (Fig.2d, Supplemental Dataset S1). More genes remain differentially expressed throughout the time course; by the end of the short time course at 4 h, mRNA abundances for 1,463 genes increased and for 1,884 genes decreased (Fig.2d). In the long time course, 9,959 genes were differentially expressed for at least one time point, with about half showing increased expression and half showing decreased expression (Fig.2d). Similar to the short time course, some genes (∼ 10%) showed a pattern of transient increase and decrease in the long time course (Fig.S2). Many genes showed changes in expression starting at various points in the time course and these were maintained throughout so that by 48 h in Fe limitation, 2,770 genes and 2,675 genes were upregulated or downregulated, respectively (Fig.2d, Supplemental Dataset S1). The overlap in differentially expressed genes (DEGs) (Fig.2c) between the short and long time courses shows that a substantial portion of the cell’s adjustment to poor Fe nutrition is already initiated by 30 min after transfer to an Fe-free environment (Fig.2d, Supplemental Dataset S1), prior to a detectable effect on cellular Fe content or photosynthesis (Fig.1c,d and Table 3), while the number of DEGs (4,681 genes) that are unique to the acclimated state indicate that there are long-term adjustments that occur between 24 and 48 h post reduction of the Fe quota.

**Table 2.**
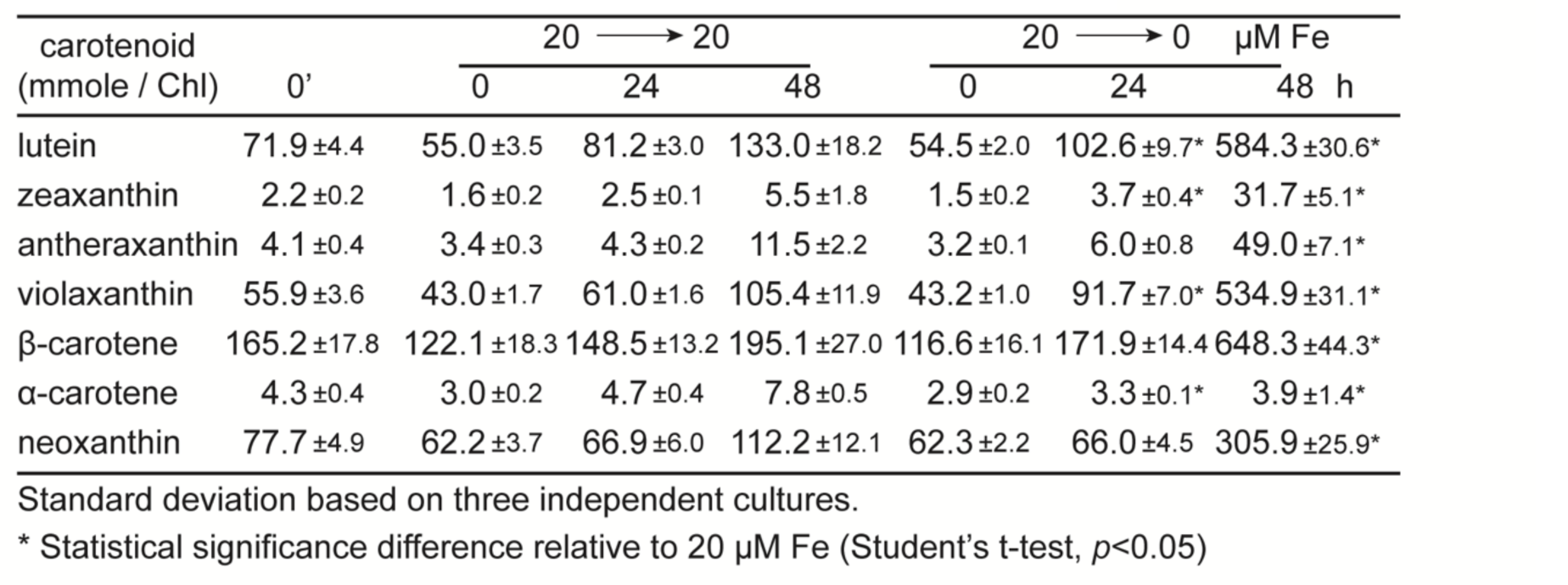
Caratenoid composition of cells transitioning into Fe limitation.

**Table 3.** Maximum quantum efficiency of PSII in cells transitioning into Fe limitation.

| time (h) | $F_v/F_m$ | |
| --- | --- | --- |
|  | 20 → 20 | 20 → 0 μM Fe |
| 0' | 0.76 ±0.01 |  |
| 0 | 0.74 ±0.01 | 0.74 ±0.01 |
| 24 | 0.75 ±0.00 | 0.70 ±0.01 * |
| 48 | 0.73 ±0.01 | 0.65 ±0.01 * |

Standard deviation based on three independent cultures.
\* Statistical significance difference relative to 20 μM Fe (Student's t-test, $p<0.05$ )

The long time course experiment showed that removal of Fe has an influence on a surprisingly large number of genes. The transcript abundance for 13,770 genes (78% of protein-coding genes in Chlamydomonas) were deemed expressed in at least one time point with a 1 FPKM minimum cutoff; of these, ∼72% showed a change in expression. By *k*-means clustering, we grouped the genes into nine clusters, each with ∼700 to ∼2700 members, based on their expression patterns (Fig.3a, Supplemental Dataset S2). Cluster 1 showed a pattern of immediate decreased transcript abundances (compare 0.5 to 0 h) upon transfer to fresh Fe-free medium and was enriched for gene ontology (GO) terms related to cilia function and assembly. These GO terms are consistent with cells in fresh medium entering G1, when genes for cilia components and biogenesis are not expressed (Zones et al. 2015; Strenkert et al. 2019). Interestingly, the peak in mRNA abundances for cluster 1 is prior to any evident physiological effect of Fe removal. Genes in clusters 2–5 all showed similar patterns of expression, with an initial increase followed by a decrease, but differed with respect to the timing of their peak mRNA abundances: early following the transfer to low Fe conditions for clusters 2 and 3, later for clusters 4 and 5. Transcript levels for genes within cluster 2 peaked early, at 30 min, and are attenuated within 2 h (Fig.3a). Protein degradation components are enriched in this cluster, consistent with previous observations of induced proteolysis in Fe-poor cells (Moseley et al. 2002; Naumann et al. 2007; Terauchi et al. 2010). It is possible that degradation of Fe-rich proteins occurs early during acclimation, as a mechanism for remobilizing Fe. Clusters 3 and 4 contained genes whose mRNA abundances peaked at 2–4 h and 8–12 h, respectively, before decreasing. These clusters are enriched for genes encoding components of anabolic metabolism related to photosynthesis and energy production. In previous work, we noted a similar pattern of increase in the abundance of Mg-protoporphyrin IX monomethylester cyclase (the di-Fe cyclase) by immunoblot analysis (Page et al. 2012). The peaks in mRNA abundances may represent stimulation of growth by transfer to a fresh medium and hence signatures of G1, which cannot be sustained in the absence of an essential nutrient, leading to the subsequent decrease by ∼24 h with RNAs in cluster 4 showing even lower abundances than at the starting point (time 0). The increase is followed by expression of components of respiration and ion homeostasis in Cluster 5, which peaked at ∼12 h. Cluster 6 was the largest cluster, containing 2,699 genes. The mRNA abundances of these genes increased late during the transition and stayed increased until the end of the experiment, likely reflecting an acclimated Fe-limited state.

**Fig.3.**
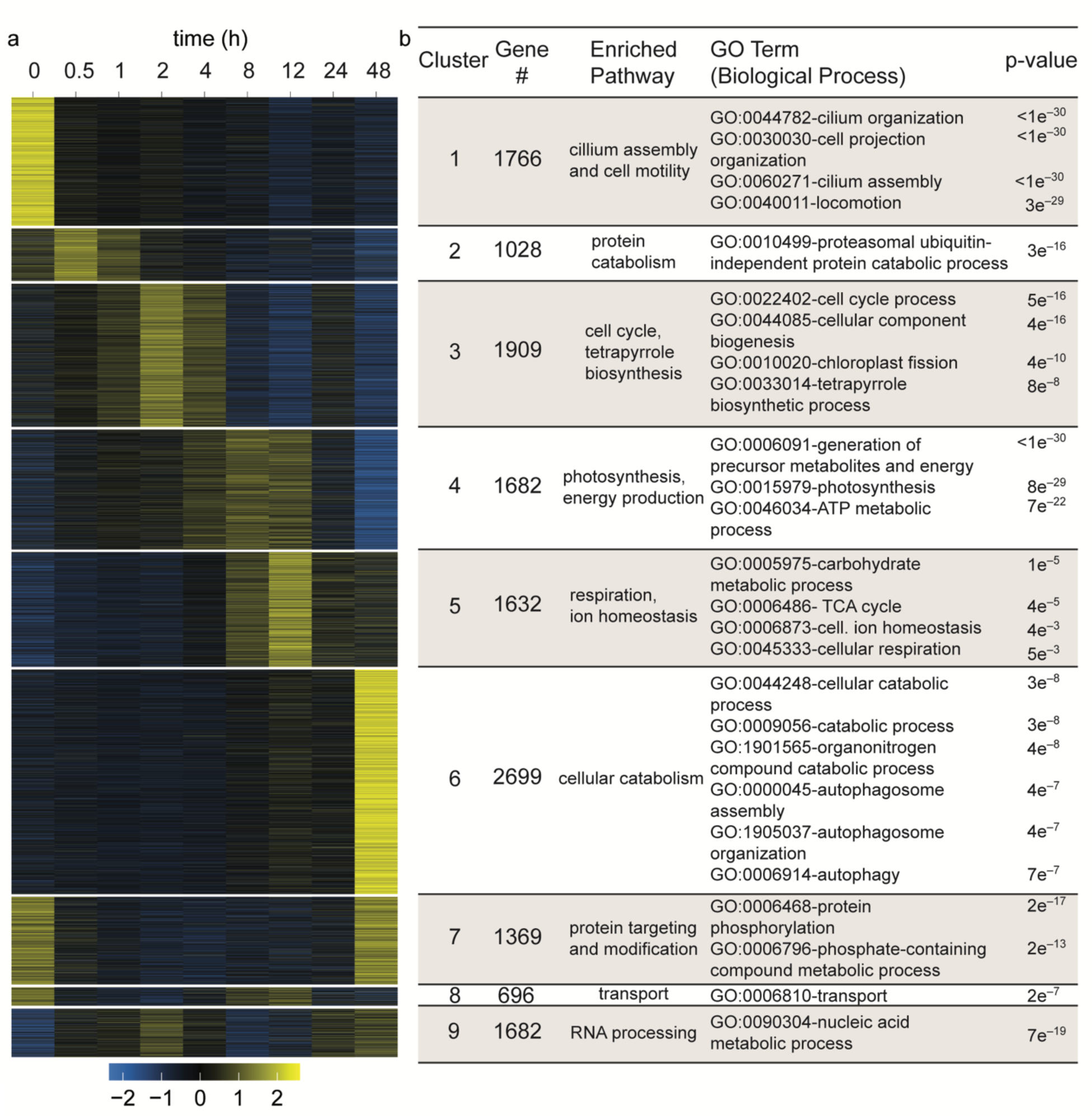
Phased response of the Chlamydomonas transcriptome following transfer to Fe-free medium. Broad, sequential changes in mRNA abundances occur over the course of 48 h in Fe-free medium. (a) Transcript abundances for nucleus-encoded genes with ≥1 FPKM in ≥1 time point (*n* = 13,770) were grouped by *k*-means clustering (*k* = 9). Here the resulting data were normalized by Z-score and plotted as a heatmap using the accompanying color scale. (b) GO enrichment analysis using the R package topGO for 9 gene clusters with the top GO terms shown (*p* value ≤ 0.05).

Clusters 7 through 9 showed unique patterns of expression with two transient peaks in mRNA abundances. For Cluster 7, we observe peak mRNA abundances in Fe-replete conditions (0 time point) with a subsequent decrease in gene expression by 30 min, and recovery by 48 h. This pattern is different from that of Cluster 1, whose constituent genes does not recover their basal expression levels at the end of the time course. Cluster 7 is enriched for genes related to protein targeting and modification. Clusters 8 and 9 contain genes whose expression pattern show two peaks during the long time course. For cluster 8, the corresponding mRNA abundances peak in Fe-replete conditions (0 time point) with an immediate decrease at 0.5 h with recovered expression at 12 h. The genes in cluster 9 peak 2 h after transfer to Fe-free medium, decrease immediately after, and recover only at the end of the time course. We conclude that the growth of Chlamydomonas under poor Fe nutrition draws on a substantial portion of the transcriptome, as cells transition through the various stages of growth (exponential to late log) and Fe nutrition (replete to limited).

### Impacts of Fe nutrition on pigments

Loss of Chl, termed chlorosis, is a signature of poor Fe nutrition. This phenotype is attributed to an Fe requirement for Chl biosynthesis (Spiller et al., 1982) and programmed degradation of the photosynthetic apparatus (Moseley et al. 2002; Terauchi et al. 2010; Yadavalli et al. 2012). Therefore, we curated the genes encoding enzymes of tetrapyrrole biosynthesis and Chl-binding proteins in more detail (Fig.4, Supplemental Dataset S3, S5). Many genes in the tetrapyrrole biosynthesis pathway are induced during the early stages of the cellular transition to Fe-limited conditions, following the cluster 3 type pattern (Fig.3a, 4a), consistent with their tight coordinate regulation in the early light phase of the cell cycle (Strenkert et al. 2019). This expression pattern perhaps reflects the stimulation of growth upon transfer of cells to fresh acetate-containing medium. Nevertheless, in the absence of Fe, the increases in mRNA abundances for genes encoding Chl biosynthesis enzymes are transient and did not result in increased Chl accumulation (Fig.4, S1). The mRNAs likely decreased eventually because the cell cannot support increased synthesis of the Fe-containing proteins in the pathway, such as the di-iron cyclase and the [2Fe-2S]-containing Chl *a* oxygenase (Tanaka et al. 1998; Tottey et al. 2003; Page et al. 2012), resulting in reduced flux through Chl biosynthesis in the Fe-poor situation. In the absence of new Chl biosynthesis, a chlorotic phenotype is established as cells grow and divide (Fig.S1, Herrin et al., 1992).

**Fig.4.**
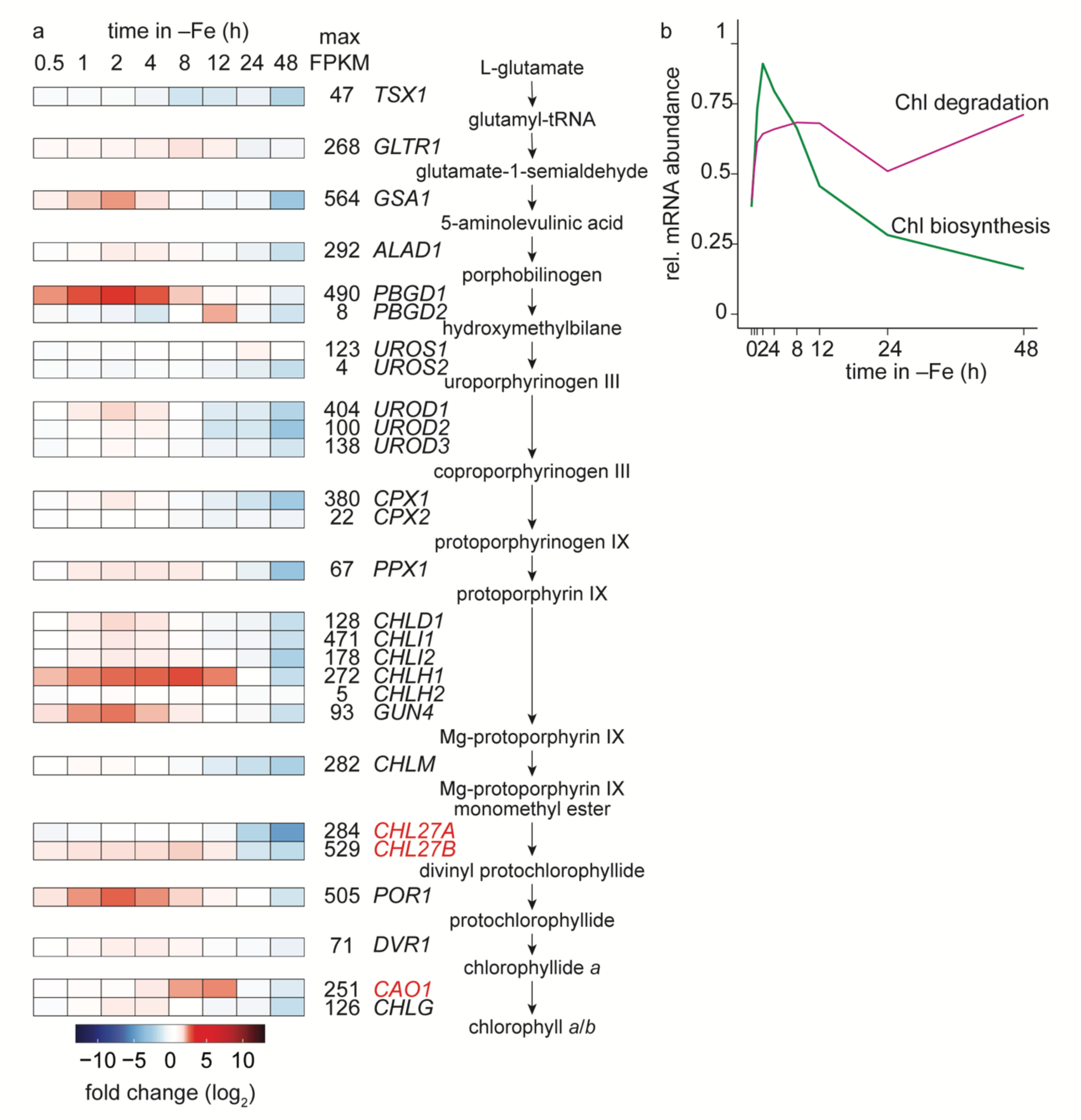
Tetrapyrrole biosynthesis genes are transiently upregulated in Fe limitation. (a) Heatmap showing mRNA abundances plotted as log_2_-transformed fold change between each time point in Fe-omitted medium relative to the 0h time point. Maximum mRNA abundances (FPKM) within all time points sampled in Fe-free medium are indicated. Arrows separate reactants and products with gene names for the corresponding enzymes indicated. mRNAs encoding Fe-binding proteins are labeled as red. Box color intensity (blue decrease; red increase) (b) Relative mRNA abundances (mRNA abundance normalized to the maximum mRNA abundance across time for each gene) averaged for all enzymes involved in Chl biosynthesis (green line) and candidate enzymes involved in Chl degradation (pink line) at the indicated time after transfer to Fe-free medium.

When we monitored the abundances of the mRNAs encoding Chl-binding proteins, we noted distinct patterns for genes encoding light-harvesting complex (LHC) proteins (*LHCA* and *LHCB/LHCBM*) vs. genes encoding other proteins like ELIPs and LHCSR3s (Supplemental Dataset S5). *LHCA*s and *LHCBs/LHCBMs* show increased expression, with mRNA abundances peaking around 8 h after transfer to Fe-free medium (cluster 4) but decaying rapidly thereafter, presumably because of feedback regulation resulting from the absence of pigment (Johanningmeier and Howell 1984) (Supplemental Dataset S5). For the other genes, *LHCSR1* parallels *LHCA*s and *LHCB*s*/LHCBMs* while *LHCSR3*s decay and are drastically less expressed by 24h (Supplemental Dataset S5). The mRNA abundances of the *ELIP*s and the *OHP*s were essentially stable during the time course (Supplemental Dataset S5). These differences may relate to the distinct functions of the various Chl-binding proteins.

A second contribution to chlorosis in Fe deficiency is from induced degradation of the photosynthetic apparatus under mixotrophy (La Fontaine et al. 2002; Moseley et al. 2002; Naumann et al. 2007; Terauchi et al. 2010; Glaesener 2019). Proteolytic degradation occurs after dissociation of LHC proteins from photosystems and occurs over long time scales (several hours). The components involved in disassembly and degradation of the proteins are not known; nevertheless, there is little change in expression of the genes encoding the putative enzymes responsible for Chl degradation (Fig.4b).

Chl-binding proteins contain carotenoids (Cars). In a situation of compromised electron transfer, as in Fe limitation, Car function may be critical for handling excess excitation energy (Yong and Lee 1991; Hagen et al. 1994, p. 3; Wang et al. 2003). Indeed, previous studies have noted that Car contents are maintained if not increased under Fe limitation (Ivanov et al. 2007; Terauchi et al. 2010; Urzica et al. 2012). Therefore, we determined Car composition over two days after transfer of cells to fresh Fe-replete or Fe-free medium (Table 2, Supplemental Table 1). We noted a net loss of Car on a per cell basis, but relative to Chl content, Cars appear retained in the Fe-free culture (Table 2, Supplemental Table 1). In the Fe-replete culture (Fe 20 → Fe 20), Car levels increased over time, reaching a maximum at 48 h, but in the Fe-free culture, the Car content increase was moderate with slightly less Car compared to the Fe-replete culture already at 24 h and dramatically less by 48 h (Supplemental Table 1). This result likely reflects an influence of poor Fe nutrition on the function of the Fe-dependent enzymes in Car biosynthesis (Fig.S3, Supplemental Dataset S4). The effect of Fe nutrition on Chl content is evident earlier in the time course (within a few hours after transfer to Fe-free medium), accounting for the higher Car/Chl already at 24 h and substantially more within 48 h (Table 2, Supplemental Table 1).

The pattern of expression of the genes encoding Car biosynthesis enzymes (Lohr 2023) is similar to that of the genes encoding enzymes of tetrapyrrole biosynthesis (cluster 3), namely a transient peak at 2 h in Fe-free medium, followed by a return to steady-state levels (Fig.S3, Supplemental Dataset S4). In general, the changes in expression of genes for Car biosynthesis are minimal, and most are downregulated by 48 h. This suggests that the overall changes in Car content in Fe-free cultures are not explained at the level of gene expression.

### Bioenergetic preference for respiration

As noted above, photosynthesis components are found in a cluster (cluster 4) different from the respiration components (cluster 5). This observation indicates different effects from poor Fe nutrition even though both pathways are dependent on Fe redox chemistry. Curation of the genes encoding individual complexes reveals distinct patterns (Fig.5). Upon transfer to fresh medium, which presumably promotes entry into G1, transcripts for genes encoding the Cyt *b*_6_*f* complex increased in abundance, and they did so before those for the photosystems (Fig.5a,b). While the *PSAs* and *LHCAs* mRNAs were coordinately expressed, the *LHCB/LHCBMs* mRNAs lagged behind *PSBs* mRNAs (Fig.5a). A similar sequential expression pattern for *PETs*, *PSAs*, *LHCAs*, *PSBs*, and *LHCB/LHCBMs* mRNAs was noted in the light phase of synchronized Chlamydomonas cells during thylakoid membrane biogenesis (Strenkert et al. 2019). After mRNA abundances peak around 12 h, they drastically decreased for all the genes encoding photosynthetic complexes except for the ATP synthase gene (Fig.5a,b vs. e), consistent with the maintenance of ATP synthase in Fe-limited cells but loss of the electron transfer complexes that are reliant on Fe (Fig.1d) (Page et al. 2012).

**Fig.5.**
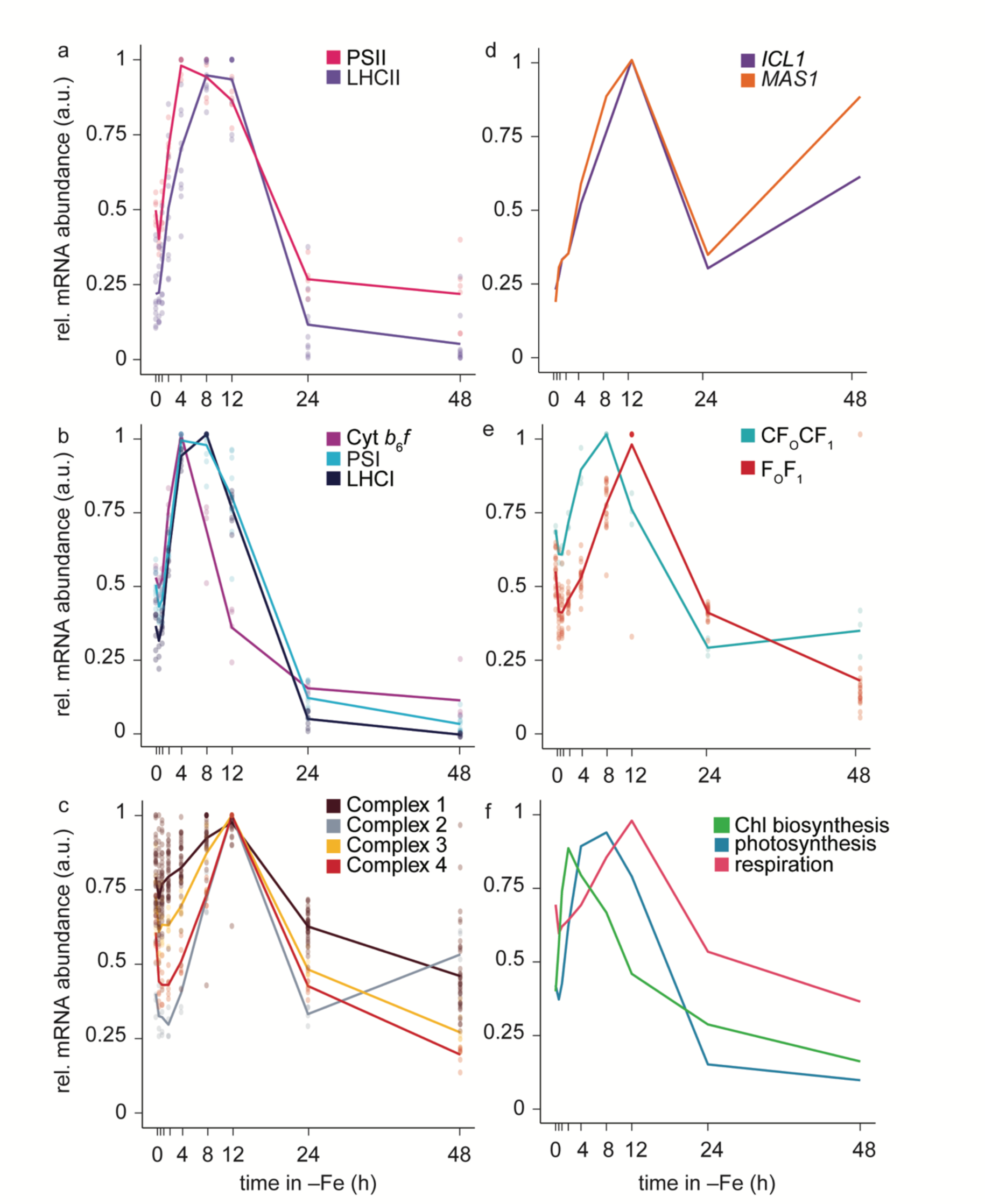
Changes in mRNAs encoding respiratory complexes are smaller than those for photosynthetic complexes. (a-e) mRNA abundances normalized to the maximum abundance across all time points. Individual subunits are plotted as points, and the average for all subunits within a complex are plotted as lines. For a complete list of genes encoding protein subunits, see Supplemental Dataset S3, S5, and S6. Photosynthesis clusters: PSII (cluster 4); LHCII (cluster 3, 4); PSI (cluster 4); LHCI (cluster 4); Cyt *b*_6_*f* (cluster 3, 4); CF_1_F_O_ (cluster 4). Respiration cluster: Complex 1 (cluster 2, 3, 4, 7), Complex 2 (cluster 5), Complex 3 (cluster 4, 8); Complex 4 (cluster 4, 8); and F_1_F_O_ (cluster 4, 6). (f) Relative mRNA abundances averaged for all genes encoding proteins in Chl biosynthesis (green), photosynthetic complexes (PSII, LHCII, PSI, LHCI, Cyt *b*_6_*f*; blue), and respiration (Complex 1-4; red).

The maximum quantum efficiency of PSII (*F_v_/F_m_*) decreased throughout the time course in Fe-free medium as cells become more Fe-starved (48 h) (Table 3), consistent with progressive loss of PSII function. Photosynthetic ferredoxin (PETF1, also reported as FDX1) is a large sink of chloroplast Fe and is a prime target for degradation in Fe-poor cells. *PETF1* and *FDX3* are co-expressed with other genes encoding components of the photosynthetic apparatus (Supplemental Dataset S5). The substrates of FDX3 are not known, but the pattern of expression suggests that the substrates may be related to the biogenesis of thylakoid membrane components. One surprising finding was the ∼45-fold change (from <1 to 31 FPKM) in *FDX2* mRNA within 12 h (Supplemental Dataset S5). FDX2 is involved in nitrate metabolism, and its synthesis is repressed by ammonium (Terauchi et al. 2010; Schmollinger et al. 2014). Perhaps this increase in the ammonium-replete Fe-poor medium is in response to FDX1 loss. FDX1 and FDX2 share substantial sequence and structural similarity that may point to overlapping function, or FDX2 may function in another pathway besides nitrate assimilation, which is activated in Fe-poor conditions (Terauchi et al. 2009). The *FDX2* response was not captured in previous work on long-term acclimation (e.g. (Urzica et al. 2012), because the mRNA levels decrease by 48 h. In agreement with previous studies, *FDX6* mRNA approximately doubled (∼54 to ∼109 FPKM) within 30 min in Fe-poor medium, and peaked by 8 h in Fe-poor medium before decreasing by 48 h, showing a similar pattern of expression to *FDX2* (Supplemental Dataset S5) (Terauchi et al. 2009).

The transcripts for genes encoding the respiratory components, Complexes I to IV and the F_1_F_o_, also showed a coordinated expression pattern (Fig.5c,e, Supplemental Dataset 6) but their abundances did not decrease in the later stages of the time course, presumably to ensure maintenance of mitochondrial energy production. We note also that transcripts encoding enzymes for acetate utilization, found in cluster 5, show a pattern like that for transcripts encoding respiratory components, but the former maintains higher levels at the later stages of Fe limitation, supporting a metabolic transition to greater reliance on heterotrophic growth (Fig.5d).

### Sentinel genes for poor Fe nutrition are expressed rapidly after transition to Fe-free medium

Previously, we noted that genes encoding components of Fe assimilation are sensitive markers of Fe status. They are upregulated in response to poor Fe nutrition even when there are no clear symptoms like chlorosis or poor growth (Fig.6a, and La Fontaine et al. 2002; Allen et al. 2007a). Of these, *FRE1*, encoding a putative ortholog of the ubiquitous eukaryotic ferrireductases that mobilize Fe(III) from chelates by reducing Fe(III) to Fe(II), was the most dramatic (Fig.6a, and Stearman et al. 1996; Robinson et al. 1997; Allen et al. 2007a). In the long time course, *FRE1* mRNA abundance increased from 0.2 FPKM to 27 FPKM during the first 30 min, and continued to increase over ∼10,000-fold to ∼2210 FPKM at 12 h (Fig.6a, Supplemental Dataset S7), becoming one of the most abundant mRNAs in the cell. In eukaryotes, mobilized Fe(II) can be assimilated either via ZIP family transporters (Eide et al. 1996; Vert et al. 2001) or via a ferroxidase–ferric transporter complex (Askwith and Kaplan 1998; La Fontaine et al. 2002; Allen et al. 2007a). Indeed, Chlamydomonas *IRT1*, *FTR1,* and *FOX1* are co-expressed with *FRE1*, albeit at different scales with *FRE1* mRNA abundance increasing ∼10,000 fold while *IRT1* (from <1 to ∼22 FPKM)*, FTR1* (∼94 to ∼1016 FPKM), and *FOX1* (∼91 to ∼802 FPKM) increased ∼8-to ∼32-fold by the first time point in the long time course and peaking within 8 to 12 h (Fig.6a, Supplemental Dataset S7). Rapid induction of *FOX1* is associated with increased FOX1 protein within 24 h of Fe limitation in photo-heterotrophic conditions (Fig.1, Fig.6a, and (Busch et al. 2008; Page et al. 2012)). The *FEA1* gene, encoding a candidate Fe-assimilation protein (Allen et al. 2007a), is expressed early at 30 min with the rest of the high-affinity pathway, while the adjacent *FEA2* paralog lags slightly behind at 1 h (Fig.6a, Supplemental Dataset S7). *IRT2* and *NRAMP4* are induced later with significant differential expression noted only at 4 h and 8 h, respectively, into the time course (Fig.6a). These results suggest a two-tiered response to poor Fe nutrition and potentially distinguish a first-line-of-defense assimilation components (*FRE1*, *FEA1*, *FOX1*, *FTR1*, *IRT1*) from Fe redistribution components (*NRAMP4*, *IRT2*) whose functions take over when assimilation becomes insufficient.

**Fig.6.**
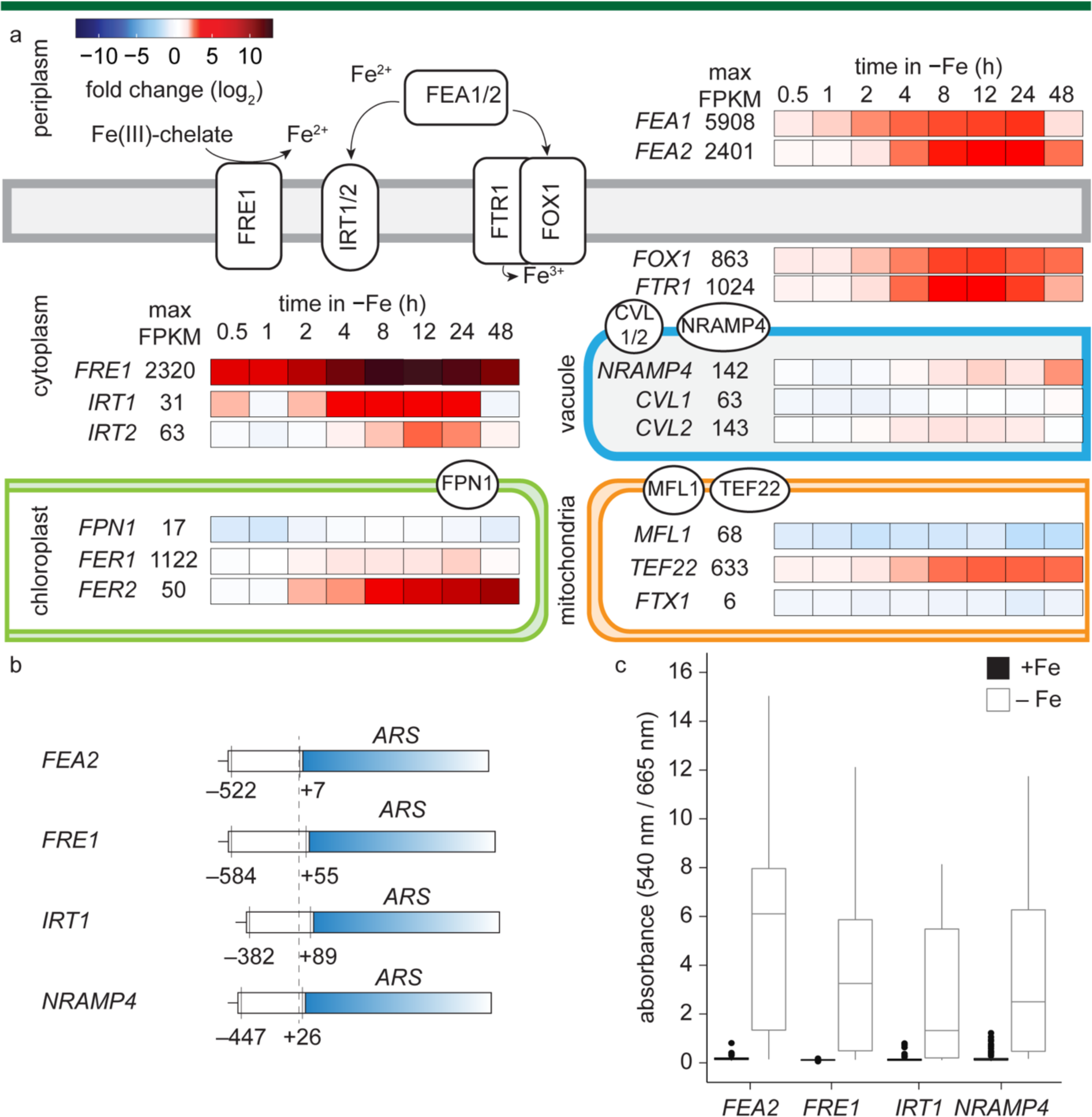
Fe assimilation pathways are transcriptionally upregulated in Fe limitation. Overview of the mRNA abundance changes involved in known and putative Fe acquisition and assimilation pathways upon Fe limitation. (a) Changes in mRNA abundances of Fe acquisition and assimilation proteins. Heatmap (red increase, blue decrease) indicates log2-transformed fold change of transcript abundance between each time point in Fe-free medium vs 0h. Presumed subcellular locations of NRAMP4, CVL1/2, MFL1, and TEF22 in Chlamydomonas are indicated. (b) *FRE1*, *FEA2*, *IRT1*, and *NRAMP4* promoter regions were fused to arylsulfatase (*ARS*) as indicated. The dashed vertical line indicates the +1 position representing the 5’ end of the corresponding transcript. (c) Each positive transformant was grown in TAP medium supplemented with either 20 μM Fe (+, black) or not (−, gray) and arylsulfatase activity was measured in the supernatant. Boxplots shows values representing an individual transformant out of a total of 96 transformants. Center line within the box indicates the median value. Top and bottom edges of the box indicate the upper and lower quartiles values, respectively. Whiskers indicate maximum and minimum data value, and points indicate outliers.

### Fe stores

Plastid ferritin and the acidocalcisome are other Fe-handling components in Chlamydomonas (Busch et al. 2008; Long et al. 2008; Blaby-Haas and Merchant 2014; Schmollinger et al. 2021; Hui et al. 2022). While ferritin is usually increased in Fe-overload situations in most organisms, in Chlamydomonas it is increased in low Fe and hypothesized to serve a role in buffering Fe released from the degradation of the photosynthetic complexes (Busch et al. 2008; Long et al. 2008). Increase in *FER1* mRNA is evident at 4 h, in a time frame compatible with the initiation of degradation of the photosynthetic complexes (Fig.6a; (Busch et al. 2008)). Ferritin1 is more abundant than ferritin2 (Busch et al. 2008; Long et al. 2008; Hsieh et al. 2012) and hence likely to be quantitatively more important in maintaining Fe homeostasis.

Fe is also stored in the acidocalcisomes (analogous to acidic vacuoles) whose boundary membranes contain CVL1 and CVL2, homologs of Arabidopsis VIT1 (Kim et al. 2006; Blaby-Haas and Merchant 2012). These transporters are likely required to re-export vacuolar Fe (Kim et al. 2006; Blaby-Haas and Merchant 2012; Long et al. 2023). *CVL2* mRNA increases within 4 h, like that of *FER1*, while *CVL1* mRNA is less abundant and not significantly increased, perhaps speaking to the greater relevance of CVL2 for maintaining Fe homeostasis (Fig.6a). *TEF22*, which is transcribed from the same promoter as *FEA1*, was identified in previous transcriptomic and proteomic experiments because of its increased expression in Fe-poor cells (Allmer et al. 2006; Urzica et al. 2012). The protein is hypothesized to function in translocation of Fe from either the chloroplast or acidocalcisomes to the mitochondria (Urzica et al. 2012). The pattern of *TEF22* expression in the long time course is similar to that of the high-affinity Fe transporters, induced within 30 min in Fe limitation and continued to increase from ∼92 to ∼633 FPKM throughout the time course (Fig.6a, Supplemental Dataset S7).

### Transcriptional regulation in response to low Fe

The rapid response of Fe-assimilation pathways to a change in medium Fe status is consistent with the involvement of transcriptional mechanisms as noted previously for *FOX1* and *FTR1* (Allen et al. 2007a). To test this idea for some of the Fe-responsive genes, we generated reporter constructs where the gene encoding arylsulfatase was placed under the control of the upstream regulatory regions of *FRE1* (577 bp), *FEA2* (529 bp), *IRT1* (471 bp), and *NRAMP4* (473 bp) and introduced them into Chlamydomonas strain CC-425 (Fig.6b). Since introduced genes insert into the Chlamydomonas genome by illegitimate recombination, the level of expression of test constructs can vary widely owing to position effects (Schroda 2019). Therefore, 96 independent transformants were assayed using a microplate-adapted version of a colorimetric assay (Blaby and Blaby-Haas 2018).

In the absence of supplemented Fe compared to cultivation in the presence of Fe, we observed on average a 31-, 30-, 17-and 21-fold increase in the activity of arylsulfatase derived from the upstream regions of *FRE1*, *FEA2*, *IRT1*, and *NRAMP4*, respectively (Fig.6c). These results suggest, as seen previously for *FOX1*, *FTR1*, and *FEA1*, that Fe assimilation and Fe mobilization are largely regulated at the transcriptional level in Chlamydomonas (Allen et al. 2007a; Deng and Eriksson 2007).

## Discussion

### Fe assimilation

Various stages of Fe nutrition in mixotrophic Chlamydomonas cells – replete, deficient, and limited – were defined based on graded phenotypes (Moseley et al., 2002). In previous work, we queried the patterns of gene expression in cells acclimated to each state, to identify 78 genes that are highly sensitive to mild decreases in Fe supply in the growth medium (Urzica et al. 2012). Most of these genes (74 of 78) also showed increased expression in cells experiencing a sustained and severe drop in Fe supply relative to Fe-replete cells (Urzica et al. 2012). In the Fe-limited situation, hundreds of other genes also exhibited changed patterns of expression relative to Fe-deficient cells, representing metabolic adjustments to the absence of an essential growth-limiting nutrient. Most of these DEGs likely represent secondary or indirect responses to poor Fe nutrition. In this work, we characterized the transition of cells from a replete condition to fresh medium lacking Fe. By monitoring the transcriptome as a function of time, we aimed to extract information on a temporal sequence of events in response to poor Fe nutrition.

The Fe content of cells in Fe-free medium decreased steadily over the 48 h period of the experiment (Fig.7a). At the start of the experiment (0 h), the Fe content matched well with the Fe content of long-term acclimated Fe-replete cultures and then progressed through a stage that matches the Fe content of cells that are long-term acclimated to Fe deficiency (Fig7a,c). Eventually, the cells reached a point (24 to 48 h) where the Fe content renders the cells growth-limited by poor Fe availability (Fig.7a,c). The temporal changes in mRNA abundances (Fig.2b) match well with the changes in the cellular Fe quota (Fig.7a): the early responses initiated at 30 min and extending up to 4 h corresponding to only small changes, followed by the later changes that initiated at 8 h and extending through 24 and 48 h. When the Fe content falls to the level measured in long-term acclimated Fe-limited cells (compare Fig.7a 24 h and 48 h points to Fig.7c 0.2 µM), the culture ceased growth and entered stationary phase. Interestingly, in an Fe-replete situation, cell transition through mild Fe deficiency during exponential growth (Fig.7b). We hypothesize that Fe-uptake, which relies on multiple redox steps, cannot keep up with the intracellular use of Fe. In previous work, we noted a transient increase in expression of Fe uptake components in such exponentially growing cultures (Page et al. 2012), and this observation illustrates the importance of the nutritional Fe regulon for cell proliferation even when external Fe supply is plentiful.

**Fig.7.**
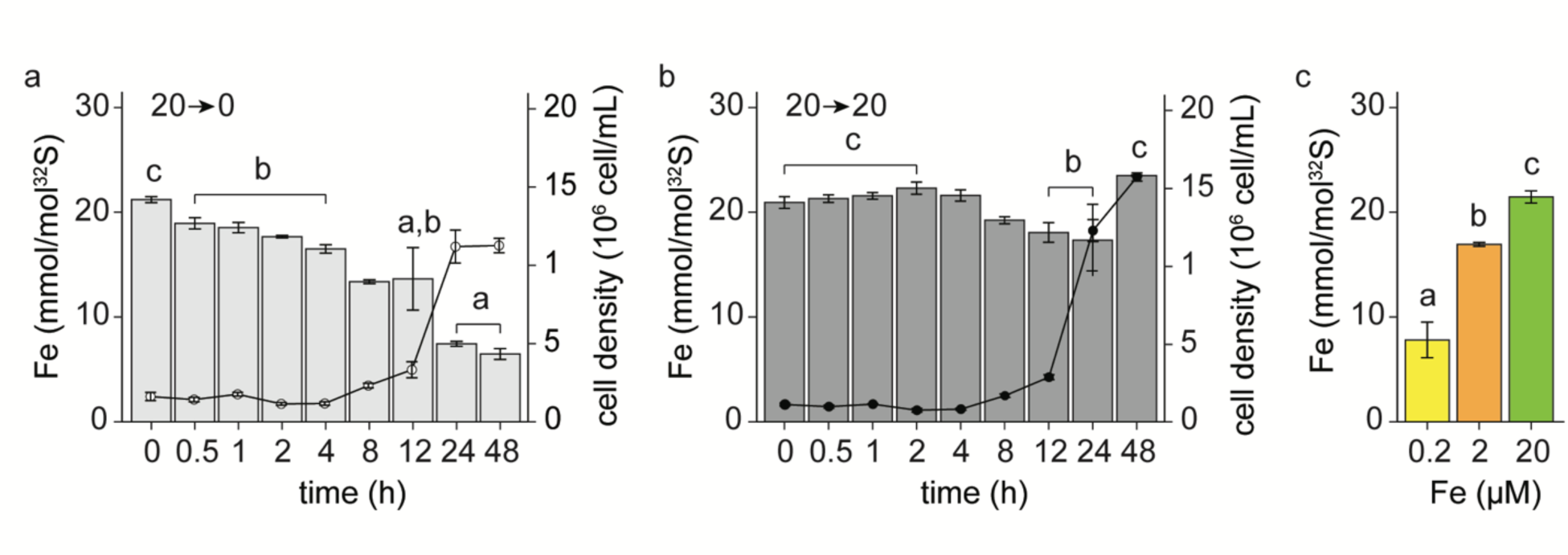
Fe nutritional stages as cells transition into Fe-free medium. (a-c) Fe content was determined by ICP-MS/MS and normalized to ^32^S content. Error bars indicate the standard error of three independent cultures. (a-b) Overlay of Fe content and cell density of cells transitioning to new medium with either (a) 0 µM Fe(light gray) or (b) 20 µM Fe (dark gray) medium. (c) Fe content at previously defined Fe nutritional stages, Fe-replete at 20 µM (green), Fe-deficient at 2 µM Fe (orange), and Fe-limited at 0.2 µM Fe (yellow). Letters (a, b, c) indicate no significant differences between cells at the corresponding sampling times compared to Fe-replete (a), Fe-deficient (b), and Fe-limited (c) steady state conditions (Student’s *t*-test, *p* > 0.05).

The Fe-assimilation pathways represent a first line of defense in the face of poor Fe nutrition. In Chlamydomonas, there are at least two likely routes for Fe uptake (Blaby-Haas and Merchant 2012): one (FOX1/FTR1) prototyped by the yeast pathway involving a multi-copper oxidase in complex with a ferric transporter, and another (IRTs) prototyped by the Arabidopsis pathway involving a ZIP family transporter (Eide et al. 1996; Vert et al. 2001). Both pathways use Fe(II) as a substrate, which is generated by a ferrireductase, an enzyme found throughout biology (Kosman 2010). We found that Chlamydomonas *FRE1* expression levels respond early and strongly to the absence of Fe in the medium (Fig.6a, Supplemental Dataset S7). The gene is generally tightly repressed in Fe-replete medium, making it a particularly sensitive marker of Fe status (Allen et al. 2007b). Although there are other candidate reductases encoded in the Chlamydomonas genome that also respond to poor Fe nutrition (Blaby-Haas and Merchant 2023), the timing and magnitude of *FRE1* expression compared to these other reductases suggests it as a key player responsible for Fe assimilation.

The FEA proteins, whose role in Fe metabolism was originally discovered in Chlamydomonas (Rubinelli et al. 2002; Allen et al. 2007a), are abundant secreted proteins. FEA-related proteins are found in many other green algae and a protein, named ISIP2a, which contains the FEA domain and shows increased expression in Fe-poor medium, was discovered recently in diatoms (Blaby-Haas and Merchant 2012; Morrissey et al. 2015; McQuaid et al. 2018). The extracellular location of FEAs suggests that they may function as substrate-binding proteins for Fe delivery to assimilation components. Indeed, one of the *FEA* genes, *FEA1*, is also expressed early in the time course like *FRE1* (Fig. 6a, Supplemental Dataset S7). The second gene, *FEA2*, likely arose by gene duplication, and although it responds within 2 h, it may have acquired additional *cis-*regulatory sequences for expression also under low inorganic carbon availability (Hanawa et al. 2007), suggesting a role for (bi)carbonate in Fe(III) binding as in the FEA-domain ISIP2a protein of diatoms and in mammalian transferrin (Fig.6a) (Lambert et al. 2005; McQuaid et al. 2018).

*IRT1*, *FOX1* and *FTR1* were other early responders. The co-expression of the corresponding proteins indicates operation of two routes for Fe uptake, but transcript abundances suggest that the FOX1/FTR1 may be the major route. While there may be some overlap, functional independence of the two possible routes is further substantiated by the Fe-nutrition-dependent growth phenotype in the *fox1* mutant (Chen et al. 2008). Each of these genes is transcriptionally regulated by Fe, as shown in this study (*IRT1*, *FRE1*, *NRAMP4*, *FEA2*) (Fig.6b) and in prior work (*FOX1*, *FTR1*, *FEA1*) (Allen et al. 2007a; Deng and Eriksson 2007). The factors mediating the transcriptional responses are not known in Chlamydomonas or other algae. We note that orthologs of *MYB10*, *PHR1*, *BTSL1/2*, and *bHLH34* involved in expression of the nutritional Fe regulon in land plants (Palmer et al. 2013; Briat et al. 2015; Vélez-Bermúdez and Schmidt 2023) are found in green algae as well (Urzica et al. 2012; Roth et al. 2017; Davidi et al. 2023) (Supplemental Dataset S8).

### Metabolism

We focused on the biosynthesis of Chl and Car. The decreased expression of Chl biosynthesis genes, occurring later in the time course (Fig.4a, Fig.S1), is a secondary response to poor Fe nutrition. Interestingly, the entire pathway is coordinately downregulated even though the key Fe-dependent step is relatively late in the pathway, suggesting that the pathway responds to a metabolic signal rather than an Fe signal (Fig.4a). The first few steps in tetrapyrrole biosynthesis are shared with heme biosynthesis and these were less affected, consistent with a continuing demand for heme by respiratory components (Fig.4a, Fig.5c, f). Many Cars are found together in Chl-binding proteins, the decrease in Chl was paralleled by the decrease in Car on a per cell basis (Supplemental Table 1). Nevertheless, the Car/Chl ratio increased over time, speaking to the differential stability of Car versus Chl, which might indicate the photoprotective roles of Car (Bassi and Dall’Osto 2021; Lohr 2023) (Table 2).

Growth inhibition from the lack of Fe is likely responsible for decreased transcript abundances after 24 h in Fe-free medium. The timing of decrease was not identical for all pathways, occurring more rapidly for genes encoding Chl biosynthesis enzymes and components of the photosynthetic apparatus than for genes encoding respiratory components (Fig.5f). This presumably reflects the cessation of transcription for the photosynthesis genes but continued expression of the genes for acetate metabolism and respiration.

### Hierarchy of nutrient limitation

Nitrogen is an essential macronutrient for plants and algae. It is a major ingredient of fertilizer and is a key limiting nutrient in marine environments. Loss of Chl and photosynthetic functions is also a notable phenotype of nitrogen (N) starvation (Martin and Goodenough 1975; Martin et al. 1976; Plumley and Schmidt 1989; Goodson et al. 2011; Schmollinger et al. 2014). A prior study of N starvation in a time course with the same Chlamydomonas strain as used in this study (Boyle et al. 2012) allowed us to make comparisons between the two situations. Principal component analysis of the two datasets showed that the bulk (40%) of the variance in the data is attributable to the lack of nutrient (N vs. Fe) with time as a second substantial (25%) contributor (Fig.8a).

**Fig.8.**
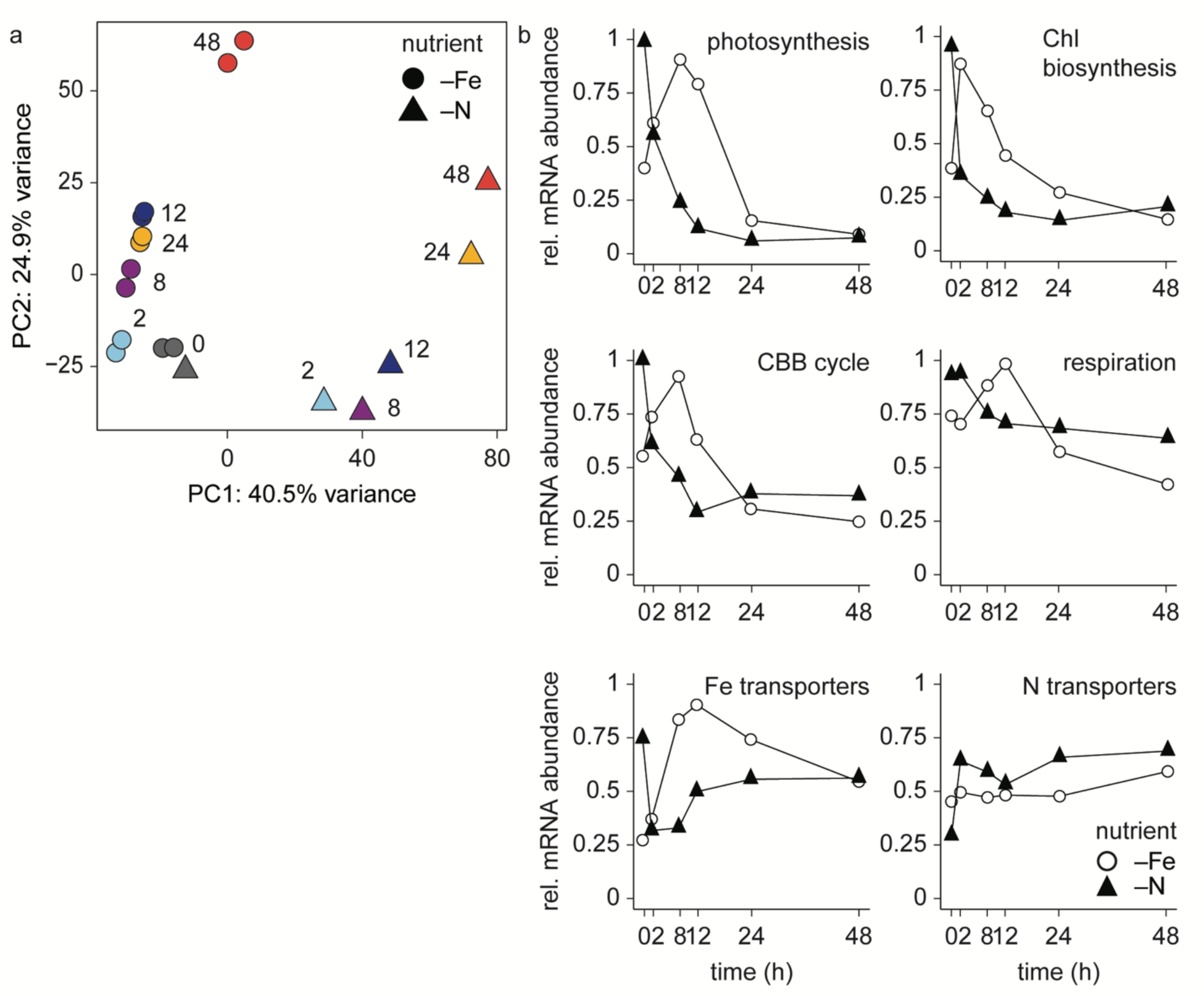
Cells under N limitation modify their transcriptome prior to cells under Fe limitation. (a) PCA of the transcriptome transitioning into Fe-(circle) or N-(triangle) limitation. Colors of each time point correspond to the length of time in the new nutrient-free medium. (b) Relative mRNA abundances (mRNA abundance normalized to the maximum mRNA abundance across time for each gene) averaged for all genes encoding enzymes involved in indicated pathways after transfer to Fe-limited (open circles) or N-limited (triangles) medium.

The 0 h time point is the most similar for both the –N and –Fe experiments, likely reflecting a common expression state at the start of the experiment: specifically, the collection, washing and transfer of replete to fresh medium (Fig.8a). When we look at the expression of genes for the photosynthetic apparatus, we see a similar decrease in transcript abundances in both cases, except there is a lag of several hours in the response to Fe starvation (Fig.8b). This likely reflects both the much higher N quota (∼10^11^/cell) (Schmollinger et al. 2014) compared to the Fe quota (∼10^8^/cell) and therefore the immediate consequences of N removal and also the existence of an Fe reservoir in the acidocalcisomes (Schmollinger et al. 2021; Hui et al. 2022; Long et al. 2023) that can buffer the effect of Fe removal. In both cases, respiration is maintained, reflecting either the smaller draw of this bioenergetic process on nutrients compared to the highly abundant photosynthetic apparatus or that acetate within the medium allows the photosynthetic apparatus to be dispensable (Fig.8b, 5c). This similarity underscores the abundance of both N and Fe held within the photosynthetic apparatus in Chlamydomonas. The pattern of expression of individual transporters is, obviously, unique for the particular nutrient: genes for N assimilation components are not induced in Fe starvation and genes for Fe assimilation components are not induced in N starvation (Fig.8b). In fact, abundances of mRNAs encoding Fe transporter are lower under N starvation, perhaps reflecting a diminished demand for Fe because of the immediate cessation of biomass production and new protein synthesis (Fig.8b).

### Summary

The present analysis provides a temporal view of the response of the Chlamydomonas transcriptome to the absence of an essential micronutrient, Fe, which includes rapid changes in expression of the assimilation pathway (*FRE1*), even before cellular Fe content is affected. This suggests the operation of a sensitive Fe-sensor, potentially one that can sense extracellular Fe bioavailability (Allen et al. 2007a), consistent with expression of reporter gene constructs (Fig.6b,c). The work also reinforces the coordinated pattern of gene expression for Chl biosynthesis and the photosynthetic apparatus noted in previous work (Duanmu et al. 2013; Strenkert et al. 2019), but now in response to a nutrient limitation (Fig.4, 5). The different consequences of poor Fe nutrition on Chl vs. Car pigments is also reminiscent of the accumulation of secondary Cars as a N deficiency response in many algae (Fig.4, Table 2) (Donkin 1976; Vechtel et al. 1992; Grung et al. 1992; Rise et al. 1994; Ben-Amotz 1995; Ho et al. 2015).

## Acknowledgments

This work was supported by Department of Energy (DOE) grant DE-SC0020627 (to SSM and SS). HWL was supported, in part, by the National Science Foundation Graduate Research Fellowship Program. Transcriptome sequencing was performed at the Broad Stem Cell Research Center High-Throughput Sequencing Core at the University of California, Los Angeles. We thank Dr. M. Dudley-Page (UCLA) for help with the promoter fusion experiments. We also thank Dr. Janette Kropat (UCLA) for ICP-MS measurements to validate the Fe status of cells used for RNA isolation. The authors declare no conflict of interest.

## Author Contributions

S.S.M., H.W.L., and S.D.G. designed the experiments and analyzed the data in the study.

E.I.U., S.D.G performed the RNA-seq experiments.

S.D.G. performed the bioinformatic analysis of RNA-seq data.

S.R.S. performed the ICP-MS experiments.

M.I. performed the pigment analysis by HPLC.

H.W.L. undertook the phenotypic analyses.

C.B. prepared and performed the reporter construct analysis.

S.S.M. and H.W.L. prepared and edited the article.

All authors commented on and revised the article.

