## Supplemental Figures and Tables for "Chlamydomonas cells transition through distinct Fe nutrition stages within 48 h of transfer to Fe-free medium"

Fig.S1 Chlorophyll content decreases upon Fe limitation
Fig.S2 Transcripts that both increase and decrease at various points throughout the time course
Fig.S3 Changes in genes encoding carotenoid biosynthesis enzymes
Supplemental Table 1. Carotenoid composition of cells transitioning into Fe limitation.

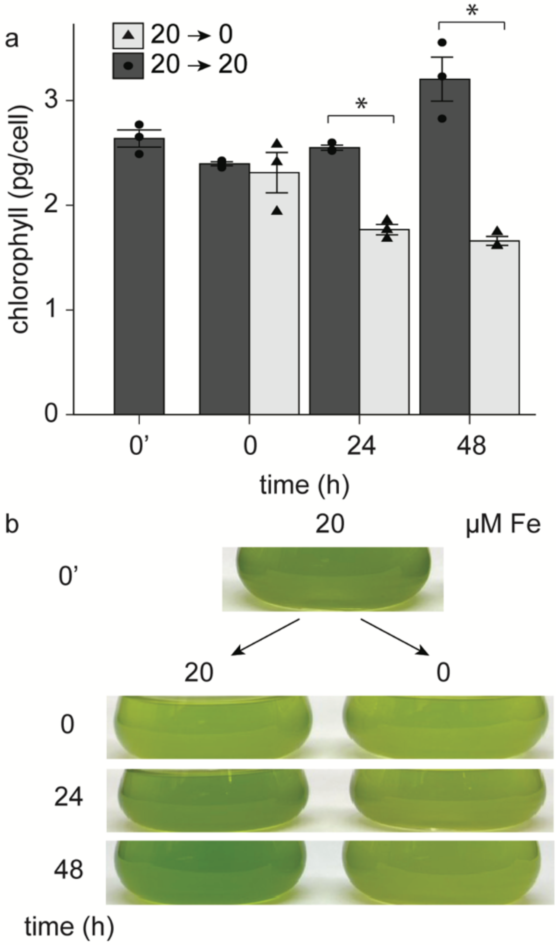


**Fig.S1 Chlorophyll content decreases upon Fe limitation**

(a) Chl content per cells in replete Fe (20 µM Fe; filled bars) or after transfer to Fe-free medium (0 µM Fe; open bars). Cells were sampled prior to medium replacement (0') or at indicated times (0, 24, 48 h). Standard error based on three independent cultures. Statistical difference relative to 20 µM for each sampling time (Student's *t*-test, $p \leq 0.05$). (b) Chlamydomonas cultures in 50 mL flasks pictured after transfer to fresh medium with the Fe concentrations as indicated. Each culture contains 2 x $10^6$ cells/mL.

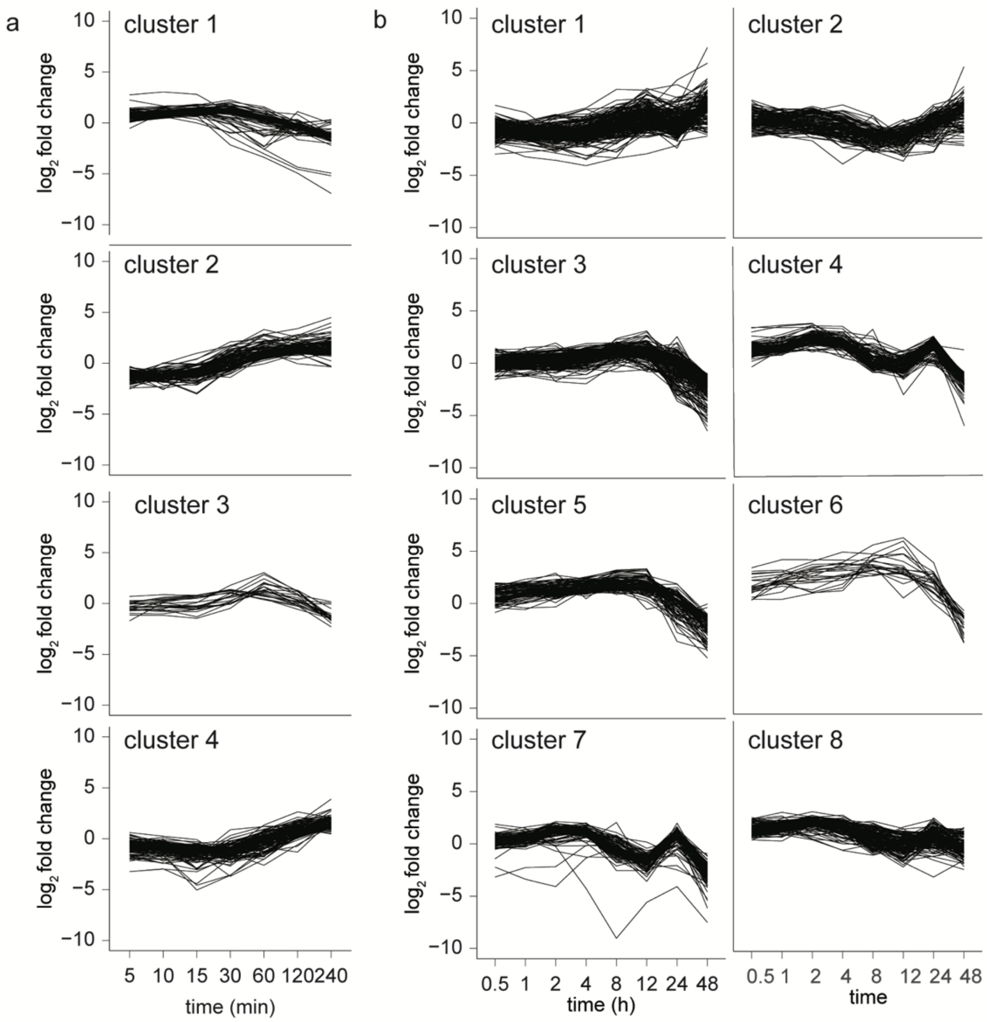


**Fig.S2 Transcripts that both increase and decrease at various points throughout the time course**

(a-b) *k*-means clustering of DEGs that reached a $\log_2$ fold change of 1 (upregulated) and −1 (downregulated). (a) The short time course includes 339 genes and was sorted into 4 *k*-means clusters, and (b) the long time course had 1,014 genes and was sorted into 8 *k*-means clusters.

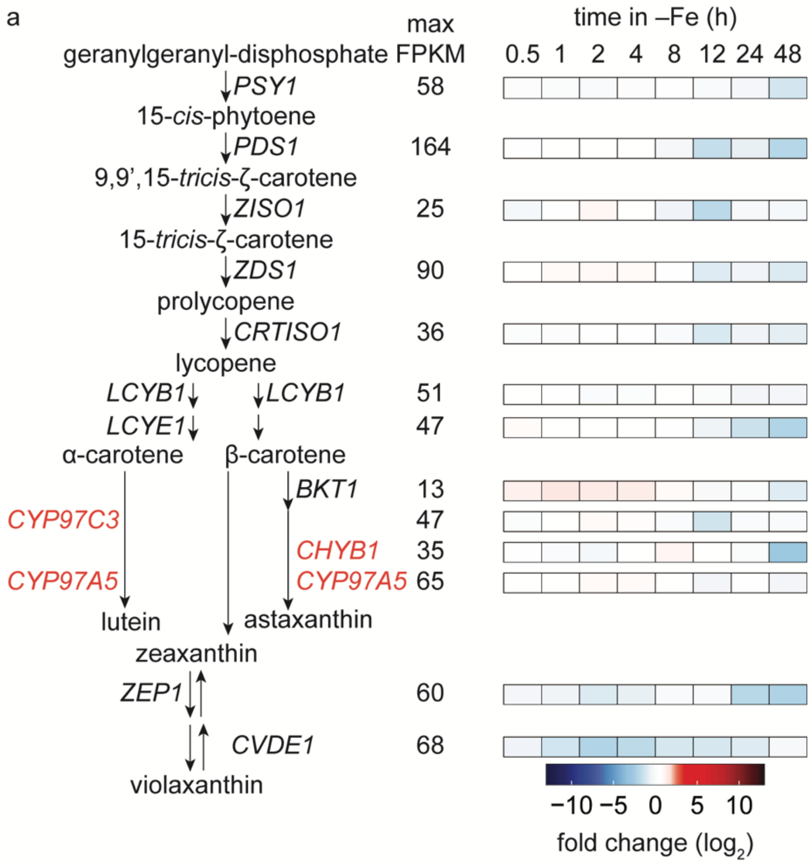


**Fig.S4 Changes in genes encoding carotenoid biosynthesis enzymes**

Arrows separate the reactants and products with the gene names corresponding to the enzymes indicated. Maximum mRNA abundances (FPKM) within all time points sampled in Fe-free medium are indicated. Heatmap (red increase, blue decrease) indicates $log_2$-transformed fold changes of sample time points vs 0 h time point. For each enzyme, only the highest expressed transcripts are shown; for a complete list of candidates, see Supplemental Dataset S4.

Supplemental Table 1. Carotenoid composition of cells transitioning into Fe limitation.

| carotenoid (attomole / cell) | | 20 ⟶ 20 | | | 20 ⟶ 0 µM Fe | | |
|---|---|---|---|---|---|---|---|
| | 0' | 0 | 24 | 48 | 0 | 24 | 48 h |
| lutein | 122.0 ±7.6 | 38.1 ±2.5 | 120.9 ±4.5 | 675.0 ±92.6 | 38.2 ±1.4 | 92.7 ±8.8* | 107.0 ±5.6* |
| zeaxanthin | 3.7 ±0.4 | 1.1 ±0.1 | 3.8 ±0.2 | 27.8 ±9.0 | 1.0 ±0.1 | 3.3 ±0.4* | 5.8 ±0.9 |
| antheraxanthin | 6.9 ±0.6 | 2.4 ±0.2 | 6.3 ±0.3 | 58.2 ±11.0 | 2.2 ±0.4 | 5.4 ±0.7* | 9.0 ±1.3* |
| violaxanthin | 95.0 ±6.1 | 29.8 ±1.2 | 90.9 ±2.4 | 535.0 ±60.8 | 30.3 ±0.7 | 82.9 ±7.1* | 98.2 ±5.7* |
| β-carotene | 281 ±30.2 | 84.6 ±12.6 | 223.0 ±19.6 | 990.0 ±137.0 | 81.7 ±11.3 | 155.0 ±13.0* | 119.0 ±8.1* |
| α-carotene | 7.3 ±0.7 | 2.1 ±0.1 | 7.0 ±0.6 | 38.9 ±3.1 | 2.0 ±0.1 | 2.9 ±0.1 | 0.7 ±0.2* |
| neoxanthin | 132.2 ±8.3 | 43.1 ±2.6 | 99.7 ±9.0 | 569.0 ±61.5 | 43.7 ±1.5 | 59.6 ±4.1* | 56.1 ±4.7* |

Standard deviation based on three independent cultures.

* Statistical significance difference relative to 20 µM Fe (Student's t-test, $p$<0.05)
